# Three-Dimensional Epigenome Roadmap of Human B-cell Differentiation Uncovers Mechanisms of Humoral Immunity and Oncogenesis

**DOI:** 10.64898/2025.12.22.695871

**Authors:** Raúl de Haro-Blázquez, Laureano Tomás-Daza, Lucía Fanlo-Escudero, Paula López-Martí, Nicolás Byrne-Álvarez, Llorenç Rovirosa, Blanca Valero-Martínez, Juan Ochoteco, Maria Rigau, Galina Medvedeva, Lucía Álvarez-González, Blanca Urmeneta, Ainoa Planas-Riverola, Jose Carbonell, Teresa Robert-Finestra, Ahlam Alqahtani, Nicholas Brittain, José Falcon-Bermejo, Marta Kulis, Jose Ignacio Martin-Subero, María Dolores Guerrero-Gilabert, Emilio Amilibia, Anna Costa, Sara Pérez, Lisa J. Rusell, Daniel Rico, Alfonso Valencia, Biola M. Javierre

**Affiliations:** Josep Carreras Leukaemia Research Institute (IJC), Badalona, Spain; Barcelona Supercomputing Center (BSC), Barcelona, Spain; Andalusian Molecular Biology and Regenerative Medicine Centre (CABIMER), Sevilla, Spain; Biosciences Institute, Newcastle University, Newcastle-upon-Tyne, UK; Institute for Biomedical Research August Pi i Sunyer (IDIBAPS), Barcelona, Spain; Biomedical Research Networking Center in Cancer (CIBERONC), Madrid, Spain; Catalan Institution for Research and Advanced Studies (ICREA), Barcelona, Spain; Barcelona University, Barcelona, Spain; Department of Otorhinolaryngology Germans Trias i Pujol Hospital (HUGTiP), Badalona, Spain; Autonomous University of Barcelona (UAB), Barcelona, Spain; Institute for Health Science Research Germans Trias i Pujol (IGTP), Barcelona, Spain

## Abstract

Human B-cell differentiation underlies humoral immunity, and its disruption leads to cancer, autoimmunity, and immunodeficiencies. However, the enhancer-based regulatory mechanisms governing this process remain poorly understood. To address this gap, we generated the first three-dimensional epigenomic roadmap of human B-cell differentiation *in vivo*. By profiling nine differentiation stages, we linked nearly half a million enhancers to their target genes through chromatin interactions, defined their activity states, and predicted bound transcription factors. We show that the acquisition of B-cell identity, distinct from common cellular processes, relies on large enhancer networks acting additively and synergistically to finely tune transcription. We further identify a 3D epigenetic priming mechanism underlying immune memory, whereby memory B cells retain primed DNA loops that enable faster responses to antigen re-exposure. Extending this roadmap to disease, we demonstrate that B-cell malignancies preserve enhancer signatures of their cell of origin while silencing distal tumor suppressor control. Finally, we uncover a previously unappreciated oncogenic mechanism in which intragenic deletions disrupt distal gene regulation via enhancer loss. Together, this resource provides a framework for understanding the genetic and epigenetic basis of humoral immunity and immune-related diseases.

## Introduction

Effective immune responses rely on the generation of a highly diverse repertoire of antigen-specific immunoglobulins (Ig), also known as antibodies or B-cell receptors, produced through B-cell differentiation^1^—a lifelong process governed by transcriptional programs that coordinate Ig diversification with cellular commitment, cell-cycle control, and metabolic reprogramming ^2^. This process unfolds in two main phases: early and late differentiation^2^. During the early stages of development, which occur first in the fetal liver and later in the fetal and adult bone marrow, hematopoietic stem cells (HSCs) give rise to uncommitted myeloid and lymphoid progenitors. The lymphoid progenitors then differentiate into early progenitor B cells—also known as PreProB cells—followed by progenitor B cells (ProB cells) and ultimately into precursor B cells (PreB cells). During these stages, B cells undergo V(D)J recombination, a genetic process that rearranges Ig gene segments to create a unique B-cell receptor. Successful cells progress to the immature B-cell state, expressing IgM on their surface, and then to the transitional B-cell state^3^. After passing negative selection to eliminate self-reactive cells, they become naive B cells, which express both IgM and IgD^4^. In the late stages of differentiation, which occurs in peripheral lymphoid organs such as the lymph nodes and spleen, naive B cells that encounter their specific antigen give rise to germinal center B cells and undergo further Ig gene diversification through class switch recombination and somatic hypermutation. These processes enhance the function and affinity of the antibody. Ultimately, germinal center B cells differentiate into either memory B cells, which confer long-term immune protection, or long-lived plasma cells, which produce and secrete large quantities of antibodies to mediate humoral immune responses^5^. Memory B cells, like naive B cells, can differentiate into germinal center B cells upon re-exposure to their specific antigen. However, they respond much faster and more robustly than naive B cells, and the mechanisms underlying this rapid immune recall response are not yet fully understood^6^.

Disruption of the specific gene transcription programs that govern B-cell differentiation can lead to immunodeficiencies, autoimmune disorders, and B-cell malignancies. Indeed, alterations in early B-cell differentiation give rise to B-cell acute lymphoblastic leukemia (B-ALL)^7^, whereas alterations affecting late B-cell differentiation primarily lead to various types of lymphomas and multiple myeloma^8,9^. Therefore, fully elucidating the molecular mechanisms that regulate transcriptional programs throughout the entire process of B-cell differentiation is crucial. Enhancers—non-coding, open chromatin regions, marked by H3K4me1 that bind transcription factors (TFs) and other regulatory proteins to activate genes through chromatin looping—stand out as key players in regulating these transcriptional programs. However, the repertoire of enhancers controlling each gene across the whole human B-cell differentiation—and their underlying regulatory mechanisms—remains unexplored due to the ethical constraints on obtaining healthy human tissue, the extreme scarcity of several B-cell populations such as progenitor or precursor B-cells, and the limited applicability of mouse data^10^. Indeed, previous studies (e.g.,^11–16^), including our own^17,18^, have identified enhancer gene associations on a limited set of B cell populations—primarily human fully committed B cells or mouse B cells derived from genetically modified knockout models that induce B-cell developmental arrest. However, a comprehensive map of human B-cell differentiation, encompassing the largely unexplored less-committed B-cell populations, is still lacking.

Here we present the first bulk, three-dimensional (3D) epigenome roadmap of human B-cell differentiation identifying the set of putative active, poised, and primed enhancers and super-enhancers, along with their bound TFs, that regulate each gene across nine key cell populations during human B-cell differentiation. In addition to CUT&RUN—profiling five histone marks—, ATAC-seq, and RNA-seq, we utilized low-input capture Hi-C (liCHi-C)^17,19^ method we previously developed to map and compare interactomes of more than 22,000 promoters at high resolution. This unprecedented resource enabled us to uncover the mechanisms by which active enhancers cooperate to drive human B-cell differentiation *in vivo* and to identify new candidate B-cell regulators. Finally, we leveraged this dataset to demonstrate its potential for clinical translation, revealing insights relevant to both adaptive immunity and oncogenesis.

## Results

### 3D Epigenome Roadmap Across Human B-cell Differentiation

To generate a gene regulatory map of human B-cell differentiation *in vivo*, we used fetal liver, fetal bone marrow, peripheral blood, and tonsils from healthy human donors to isolate nine key B-cell populations representing distinct stages of differentiation (Fig. 1A; Extended Data Fig. 1A; Methods). These included hematopoietic stem cells (HSC), early progenitor B cells—also known as PreProB cells—, progenitor B-cells (ProB), precursor B-cells (PreB), a mixed population of immature and transitional B-cells (immtransB), naive B cells (nB), germinal center B cells (GCB), memory B cells (memB), and plasma cells (PC). In addition, we isolated three control hematopoietic cell types: common myeloid progenitors (CMP) and monocytes (Mon) as controls from the myeloid lineage, and naive CD8⁺ T cells (nCD8) as a control from the T lymphoid lineage (Extended Data Fig. 1A-B). These cell populations were analyzed using CUT&RUN profiling of five histone marks—H3K4me3, H3K4me1, H3K27ac, H3K27me3, and H3K9me3—along with ATAC-seq to identify enhancers and their activities—active, primed or poised— as well as the predicted bound TFs (Extended Data Fig. 1C; Supplementary Table 1). We also analyzed these cell populations using the liCHi-C method to associate enhancers with candidate target genes (Extended Data Fig. 1C; Supplementary Table 1), whose expression levels were quantified by RNA-seq (Extended Data Fig. 1C; Supplementary Table 1). Collectively, this extensive resource—comprising 356 valid datasets across up to five highly reproducible biological replicates per omic modality and cell type in primary human samples, with a total of more than 5.5 TB of raw data—allowed us to identify and assign 468,651 high-confidence enhancers (covering 562 Mb) to their candidate target genes, as well as to predict their activity levels and associated TFs (Extended Data Fig. 2; Supplementary Table 2).

**Figure 1.**
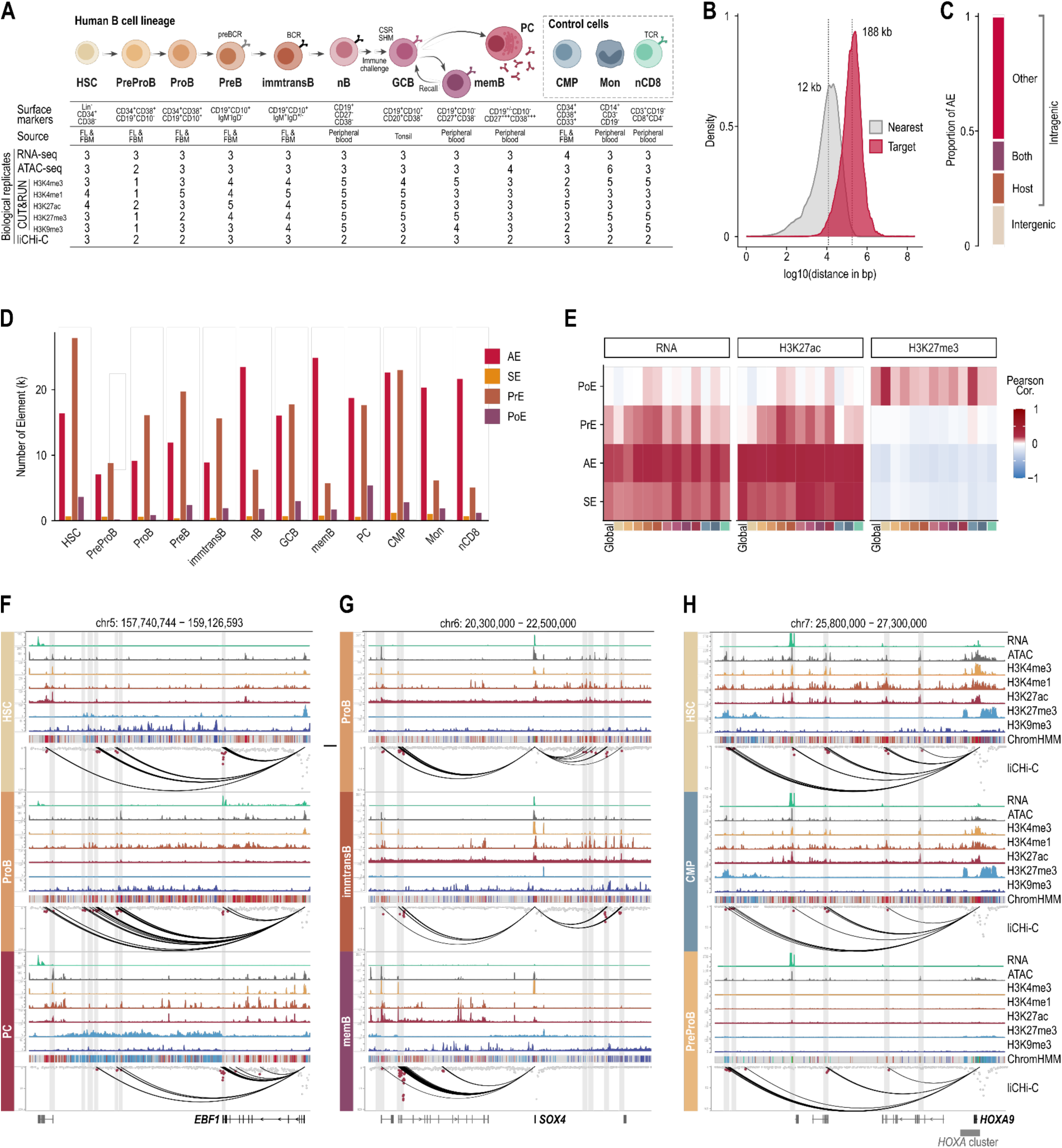
3D Epigenome Roadmap Across Human B-cell Differentiation. **A.** Experimental multi-omics framework of the work. **B.** Density distribution of genomic distances between regulatory elements and transcription start sites (TSSs). Grey indicates the distance to the nearest protein-coding gene TSS, whereas red indicates the distance to the promoter of the target gene assigned through significant liCHi-C interactions. Vertical dashed lines denote median distances for each distribution. **C.** Genomic annotation of active enhancers (AEs). Bars represent the proportion of AEs located in intergenic regions or within gene bodies (intragenic). Intragenic enhancers are further classified according to whether their liCHi-C–assigned target corresponds to the host gene, a different gene, or both. **D.** Number of regulatory elements identified per cell type, stratified by chromatin state: active enhancers (AE), super-enhancers (SE), primed enhancers (PrE), and poised enhancers (PoE). **E.** Heatmap summarizing Pearson correlation coefficients between the number of promoter-linked regulatory elements per gene and (left) gene expression levels, (middle) promoter H3K27ac signal, and (right) promoter H3K27me3 signal. Correlations are shown globally and stratified by regulatory element class. Color scale indicates correlation strength and direction. **F.** *EBF1* regulatory landscape in hematopoietic stem cells (HSC, top), progenitor B cells (ProB, middle), and plasma cells (PC, bottom) according to RNA, open chromatin, histone modifications, ChromHMM states and liCHi-C data. **G.** *SOX4* regulatory landscape in progenitor B cells (ProB, top), immature-transitional B cells (immtransB, middle), memory B cells (memB, bottom), according to RNA, open chromatin, histone modifications, ChromHMM states and liCHi-C data. **H.** *HOXA9* regulatory landscape in hematopoietic stem cells (HSC, top), common myeloid progenitors (CMP, middle), and PreProB cells (bottom) according to RNA, open chromatin, histone modifications, ChromHMM states and liCHi-C data.

Specifically, we associated an average of 16,745 active enhancers, 14,261 primed enhancers, 2,222 poised enhancers, and 689 super-enhancers to candidate target genes per cell population (Fig. 1D). Remarkably, we found that enhancers interact with their target genes over a median linear distance of 188 kb (SD = 26 kb), which is 15 times greater than the median distance to the nearest gene (12 kb; SD = 1 kb) (Fig. 1B). Only 12% of enhancers were linked to the closest gene, with some enhancers located more than 1 Mb away from their targets, such as in the case of the *EBF1, SOX4* and *HOXA9* (Fig. 1F-H; Extended Data Fig. 3). Interestingly, 81% of enhancers were located within gene bodies; however, only 33% of these intragenic enhancers were associated with their host gene, while 91% were linked to other genes, as exemplified by the regulation of *SOX4* (Fig. 1C,G; Extended Data Fig. 3B). These findings underscore the importance of mapping long-range interactions between promoters and regulatory elements to avoid misleading gene associations based on proximity.

We next validated enhancer–target gene associations by integrating RNA-seq data (Fig. 1E). A positive correlation was observed between the number of active enhancers or super-enhancers and the expression levels of their candidate target genes. This correlation was weaker for primed enhancers and markedly lower for poised enhancers. These trends were further supported by H3K27ac levels at the promoter regions (−1000+200bp from gene TSS) of the predicted target genes, used as a proxy for transcriptional activity. An opposite trend was observed for H3K27me3 levels at the same promoter regions—a histone modification associated with gene silencing.

To further support this validation, we visually examined the regulatory landscapes of genes with well-established roles in B-cell differentiation. For instance, *EBF1*—a gene encoding a key TF for B-cell lineage commitment, whose loss causes lymphopenia and immunodeficiency, while mutations are linked to B-ALL— is lowly expressed in HSCs but strongly induced in progenitor B cells and maintained until plasma cells, where it declines (Fig. 1F; Extended Data Fig. 3A). Our data suggest that transcription is regulated by dynamic enhancers located in seven distal regulatory regions (DRR1-7, from nearest to farthest) between 1.27 Mb and 0.25 Mb downstream of the transcription start site. *EBF1* expression during early B-cell differentiation, such as in progenitor B cells, is likely strongly supported by active enhancers in DRR2-7, as well as a super-enhancer in DRR1. Conversely, in HSC, CMP, plasma cells, monocytes, and nCD8 cells —where *EBF1* expression is very low—most of these DRRs are characterized as primed enhancers or non-enhancers. Similarly, SOX4 is a double-edged TF since it is essential for progenitor B and immature and transitional B-cells cell survival and differentiation, but pathogenic when dysregulated—contributing to immunodeficiency when underexpressed, leukemia and lymphoma when overexpressed, and potentially autoimmunity when misregulated. In progenitor B cells, the high *SOX4* expression is potentially supported by active enhancers in DRRs spanning a 2 Mb region, whereas in immature and transitional B cells it is supported by active enhancers in five or these DRRs. Contrarily, in memory B cells—where *SOX4* expression is very low—most of these regions are not identified as enhancers (Fig. 1G; Extended Data Fig. 3B). Contrarily, *HOXA9* encodes a TF highly expressed in HSCs, promoting their self-renewal and proliferation (Fig. 1H; Extended Data Fig. 3C). Its expression decreases during B-lineage commitment, enabling activation of key transcription factors such as *EBF1* and *PAX5* that drive B-cell differentiation. According to our data, *HOXA9* expression is potentially controlled by six DRRs between 1.34 Mb and 0.32 Mb downstream of the transcription start site. In HSC, *HOXA9* expression—unlike in the other committed cell populations such as CMP—is likely to be primarily driven by active enhancers located in DRR5, DRR3, and DRR1, as well as by super-enhancers within DRR4 and DRR2. Contrarily, DRR6 is enriched in poised enhancer states.

Collectively, these findings, supported by many additional examples (Extended Data Fig. 4-7), validate the accuracy and functional relevance of the enhancer–target gene assignments and highlight the overall quality and potential of the 3D epigenome roadmap for uncovering novel mechanisms driving human B-cell differentiation and B-cell related diseases.

### Dynamic Enhancer-Promoter Associations Shape Transcriptional Decisions During B-cell Differentiation

Given their recognized role in gene transcription, we focused on active enhancers—defined as open chromatin regions marked by H3K4me1 and H3K27ac—that were connected to distal target genes through significant liCHi-C interactions. Principal component analysis (PCA) of H3K27ac signals across these genomic regions identified as active enhancers clearly separates the different cell populations, confirming the robustness and specificity of the active enhancer identification (Extended Data Fig. 8A). Hierarchical clustering of promoter interactions with genomic regions harboring an active enhancer identified in any cell population showed a robust organization that mirrors the B-cell differentiation trajectory and the broader hematopoietic lineage hierarchy (Fig. 2A). Clustering of these interactions identified cell type-specific clusters (e.g., clusters 20, 22-24) as well as early and late B differentiation-specific clusters (e.g., clusters 25,10) (Fig. 2A; Extended Data Fig. 8B). These results not only validate the quality of gene–target associations inferred from 3D chromatin topology but also demonstrate that dynamic, cell type- and lineage-specific promoter–active enhancer interactions coordinate transcriptional programs during B-cell differentiation.

**Figure 2.**
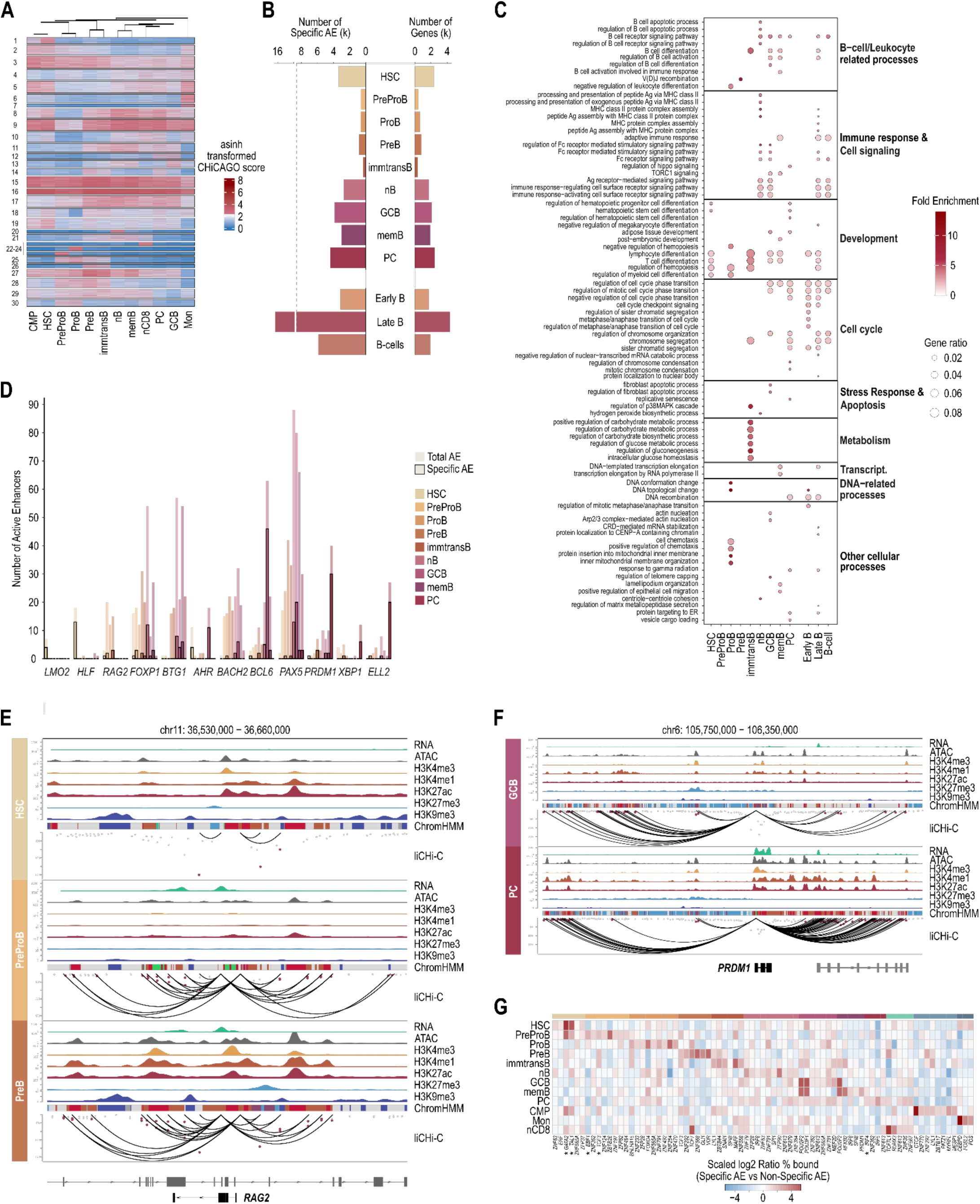
Dynamic Reconfiguration of Active Enhancer–Promoter Contacts Guides Transcriptional Programs During B-Cell Differentiation. **A.** Heatmap of asinh-transformed CHiCAGO scores for significant promoter–active enhancer (AE) interactions detected in at least one cell type. Rows correspond to AE clusters identified by k-means clustering, and columns correspond to cell populations. Hierarchical clustering of cell types (top dendrogram) was performed using average linkage and Euclidean distance. Heatmap colors represent CHiCAGO interaction strength after transformation (blue, not significant, CS < 5; red, significant). **B.** Number of cell type–specific active enhancers (left) and their associated target genes (right) identified in each individual cell population and in aggregated developmental groups (Early B, Late B, and B cells). **C.** Functional enrichment analysis of genes linked to cell type–specific AEs. Dot plots show significant Gene Ontology biological process terms grouped by major functional categories. Dot size indicates gene ratio, and color denotes fold enrichment. **D.** Number of total active enhancers (AE) (filled bars) and the subset classified as cell type–specific AE (outlined bars) across all profiled cell populations, shown for selected transcription factors. **E.** *RAG2* regulatory landscape in hematopoietic stem cells (HSC, top), PreProB cells (middle), and precursor B cells (PreB, bottom) according to RNA, open chromatin, histone modifications, ChromHMM states and liCHi-C data. **F.** *PRDM1* regulatory landscape in germinal center B cells (GCB, top), and plasma cells (PC, bottom) according to RNA, open chromatin, histone modifications, ChromHMM states and liCHi-C data. **G.** Heatmap of the ratio of transcription factor (TF) binding in cell type-specific vs. non-specific active enhancers. The top 10 TFs most enriched in cell type-specific enhancers (identified from per cell-type residual analysis, see methods) are shown, collapsed to motif cluster representatives. Colors at the top correspond to the cell type in which the TF was identified (ordered as in the rows). Heatmap colors correspond to relative enrichment (blue = low, red = high) after scaling per row.

To examine these observations in greater detail, we defined the set of cell population-specific active enhancers and their target genes, as well as those uniquely associated with early B cells (i.e., conserved in at least 50% of populations from PreProB through immature and transitional B cells), late B cells (i.e., conserved in at least 50% of populations from naive B cells through plasma cells and memory B cells), and broadly across all B-cell populations (i.e., conserved in at least 5 out of the 8 B-cell populations) (Fig. 2B; Supplementary Table 2). Focusing on genes with well-established roles in B-cell differentiation and function, we found that both the number of specific active enhancers and the overall number of active enhancers correlated closely with the cell populations in which these genes are known to function (Fig. 2D). For instance, TFs recognized as master regulators of stem cell potential, such as *LMO2* and *HLF*, were primarily associated with specific active enhancers in HSCs (Extended Data Fig. 4A-B). In contrast, *RAG2* was predominantly linked to specific active enhancers in progenitor and precursor B cells, where V(D)J recombination takes place (Fig. 2E; Extended Data Fig. 4C). Additionally, *BCL6*, *AHR*, and *BACH2* showed associations with specific active enhancers in germinal center B cells, while *XBP1*, *ELL2*, and *PRDM1* were primarily found at specific active enhancers in plasma cells (Fig. 2F, Extended Data Fig. 5-6). Furthermore, *FOXP1—* encoding a transcriptional repressor—and *BTG1* antiproliferative gene were most strongly associated with specific active enhancers in naive and memory B cells, respectively (Extended Data Fig. 7).

Additionally, we performed a gene-agnostic analysis using Gene Ontology (GO) that identified these specific active enhancers as master regulators of biological processes involved in B-cell development, B-cell function and immune response (Fig. 2C). For example, active enhancers specific to precursor B cells regulate Ig diversification via V(D)J recombination—a defining feature of this cell population. Furthermore, these specific active enhancers control cell cycle, stress response, apoptosis and metabolism—key processes dynamically modulated during B-cell differentiation. Finally, we compiled a manually curated list of genes relevant to human B-cell differentiation by integrating data from B cell–related GO terms and literature with our RNA-seq results (Extended Data Fig. 8C; Supplementary Table 3).

To gain insight into the mechanisms underlying cell type–specific enhancer activity, we identified putative TFs occupying these regions. The most enriched TFs showed markedly dynamic binding patterns across B-cell differentiation at these cell type-specific active enhancers, encompassing known regulators of specific B-cell subsets as well as TFs whose roles in B-cell biology remain uncharacterized (Fig. 2G; Extended Data Fig. 9). For instance, GATA2 and TAL1 were among the top ten TFs predominantly bound to specific active enhancers of HSCs. In contrast, EBF1, TCF3 and IRF4 ranked among the top ten TFs predominantly bound to specific active enhancers of early progenitor B cells, precursor B cell and plasma cells, respectively (Fig. 2G; Extended Data Fig. 9). This dynamic enrichment pattern indicates that cell type–specific active enhancers act as integrating platforms of sequential and population-specific regulatory inputs from environmental and developmental cues through TF binding, thereby orchestrating the transcriptional program underlying B-cell fate determination.

Collectively, these findings reveal that active enhancer–target gene interactions are dynamically and coordinately remodeled throughout B-cell differentiation, integrating diverse regulatory cues to sculpt the transcriptional programs underlying B-cell identity and function.

### Cell Type–Specific Active Enhancers Reveal New Candidate Regulators of B-Cell Identity

In addition to well-characterized genes and processes, our analysis of cell population–specific active enhancers paves the way for identifying previously unrecognized genes essential for human B-cell differentiation and function. Given their recognized role in cell fate, we focused our analysis on TFs regulated by B-cell-specific active enhancers, whose dynamic expression across B-cell differentiation correlated with the number of active enhancers. Among these TFs, MIXL1, which has no previously reported link to B-cell differentiation or function, emerges as a novel candidate TF driving germinal center B-cell biology. MIXL1 regulates transcriptional networks involved in cell cycle control, differentiation, and self-renewal. It has been shown to influence embryonic development, particularly mesodermal differentiation, as well as early hematopoiesis and stem cell potential^20^, but it has not been directly involved in the regulation of mature immune cells, such as germinal center B cells. In addition, it has been implicated in cancer biology, particularly in hematological malignancies such as leukemia^21,22^. Our data show that *MIXL1* is selectively expressed at both the mRNA and protein levels in germinal center B cells, with low expression in plasma cells and no detectable expression in other B-cell populations (Fig. 3A,C-D). This expression is regulated by six active enhancers, four of which are germinal center–specific and two shared with plasma cells (Fig. 3A-C). Notably, footprinting analysis identified MIXL1 binding at 1,820 TF binding sites (TFBS) in germinal center B cells (Fig. 3E-F), supporting a pioneering role for this TF in germinal center B-cell biology.

**Figure 3.**
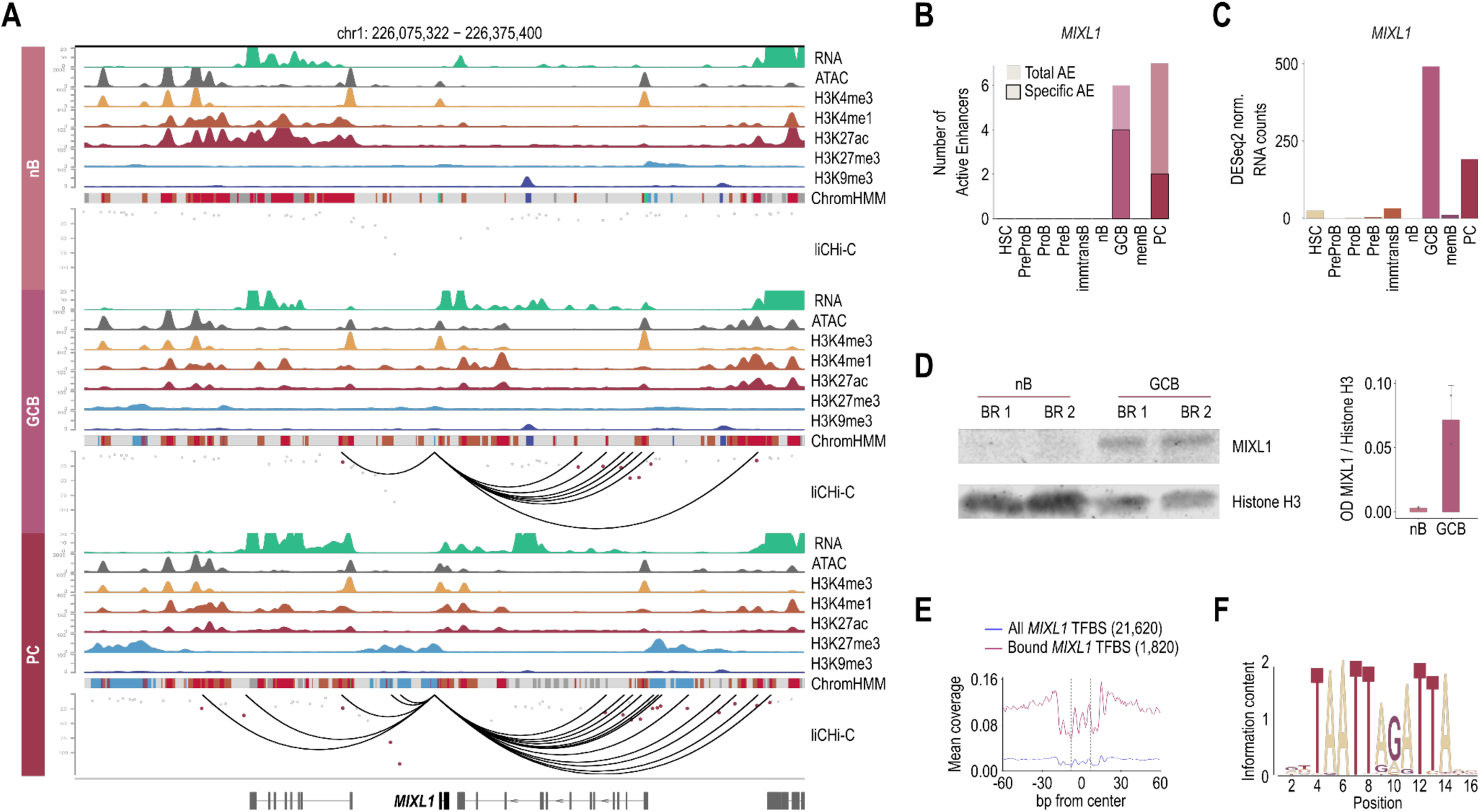
Cell–Specific Active Enhancers Points to Previously Unrecognized Identity Regulators. **A.** *MIXL1* regulatory landscape in naïve B cells (nB, top), germinal center B cells (GCB, middle), and plasma cells (PC, bottom) according to RNA, open chromatin, histone modifications, ChromHMM states and liCHi-C data. **B.** Total number of active enhancers (AE) (filled bars) and the subset classified as cell type–specific AE (outlined bars) across all profiled cell populations for *MIXL1*. **C.** Gene expression (DESeq2 normalized counts) of *MIXL1* across all profiled cell populations. **D.** Western blot of nuclear fraction of MIXL1, using Histone 3 (H3) as control (left) and associated quantification (right). **E.** Aggregate transcription factor footprinting analysis for *MIXL1* in germinal center B cells. The blue profile represents the average footprint across all *MIXL1* motif instances overlapping open chromatin regions, whereas the red profile corresponds to the subset of transcription factor binding sites (TFBS) predicted to be bound by MIXL1. **F.** DNA sequence logo of the *MIXL1* binding motif derived from predicted bound sites. The total height of each position reflects information content relative to a uniform nucleotide distribution, and the height of individual bases represents their relative frequency at that position.

Furthermore, we also identified TFs that have been suggested in a few studies to participate in B-cell biology, although their roles remain incompletely defined due to the lack of *in vivo* validation in humans. Among these is the *MYBL1* gene, also known as *MYB proto-oncogene-like 1*. It encodes a TF that regulates cell proliferation and has been primarily implicated in spermatogenesis, mammary gland development, and various cancers—including B-cell lymphomagenesis^23,24^. More recently, it has been proposed to be required for the enhanced *in vitro* differentiation of antibody-secreting cells through inhibition of the Polycomb repressive complex 2^25^. According to our data, *MYBL1* is expressed at low levels throughout human B-cell differentiation, except in germinal center B cells, where its expression is approximately 20-fold higher, driven by 11 active enhancers—6 of which are specific to this B-cell population (Extended Data Fig. 10A-C). Globally, these active enhancers are already active in naive B cells but cannot drive gene transcription because they are not in proximity to the *MYBL1* promoter. A similar regulatory landscape is observed in memory B cells (data not shown), whereas in plasma cells these enhancers are found in a primed state and remain physically distant from the *MYBL1* promoter, thus failing to induce transcription in these B-cell populations. In addition, 3,634 binding events of MYBL1 were identified in germinal center B cells, demonstrating the functional activity of MYBL1 (Extended Data Fig. 10D-E).

Although functional validation— which remains particularly challenging due to the limited capacity to expand and genetically manipulate primary human B cells and the substantial interspecies differences— is required to fully establish the roles of MIXL1, MYBL1, and other TFs identified in this analysis, these results underscore the value of 3D epigenome profiling in uncovering novel regulators of cell fate and function.

### Additive and Synergistic Active Enhancer Networks as Key Regulators of B-cell Fate and Function

We then focused our analysis on understanding the mechanistic principles by which active enhancers regulate B-cell fate. We observed an average of 3.7 active enhancers regulating each gene per cell type (Fig. 4A). To further characterize this architecture of multi-enhancer–gene associations, we constructed gene-centric active enhancer networks, representing the collection of active enhancers putatively regulating each gene in a cell population. These active enhancer networks spanned a mean size of 752 kb, underscoring the importance of 3D chromatin organization in long-range regulation and setting them apart from super-enhancers (Extended Data Fig. 11A). The analysis of the degree of centrality revealed that B-cell–specific genes—such as TFs (e.g., *IKZF1*, *PAX5*, *BCL6*, *ETS1*), proteins involved in V(D)J recombination and class switch recombination (e.g., *HMGB1*, *ZMYND8*), and regulators of B-cell receptor signaling (e.g., *LYN*, *VAV3*)—are among the most densely regulated by active enhancers, often in a highly dynamic and cell type–specific manner (Fig. 4B-D; Extended Data Fig. 11B). This was further supported by the observation that genes relevant to human B-cell biology were regulated by an average of 6 active enhancers, 2 times more than non-B-cell genes (Extended Data Fig. 11C).

**Figure 4.**
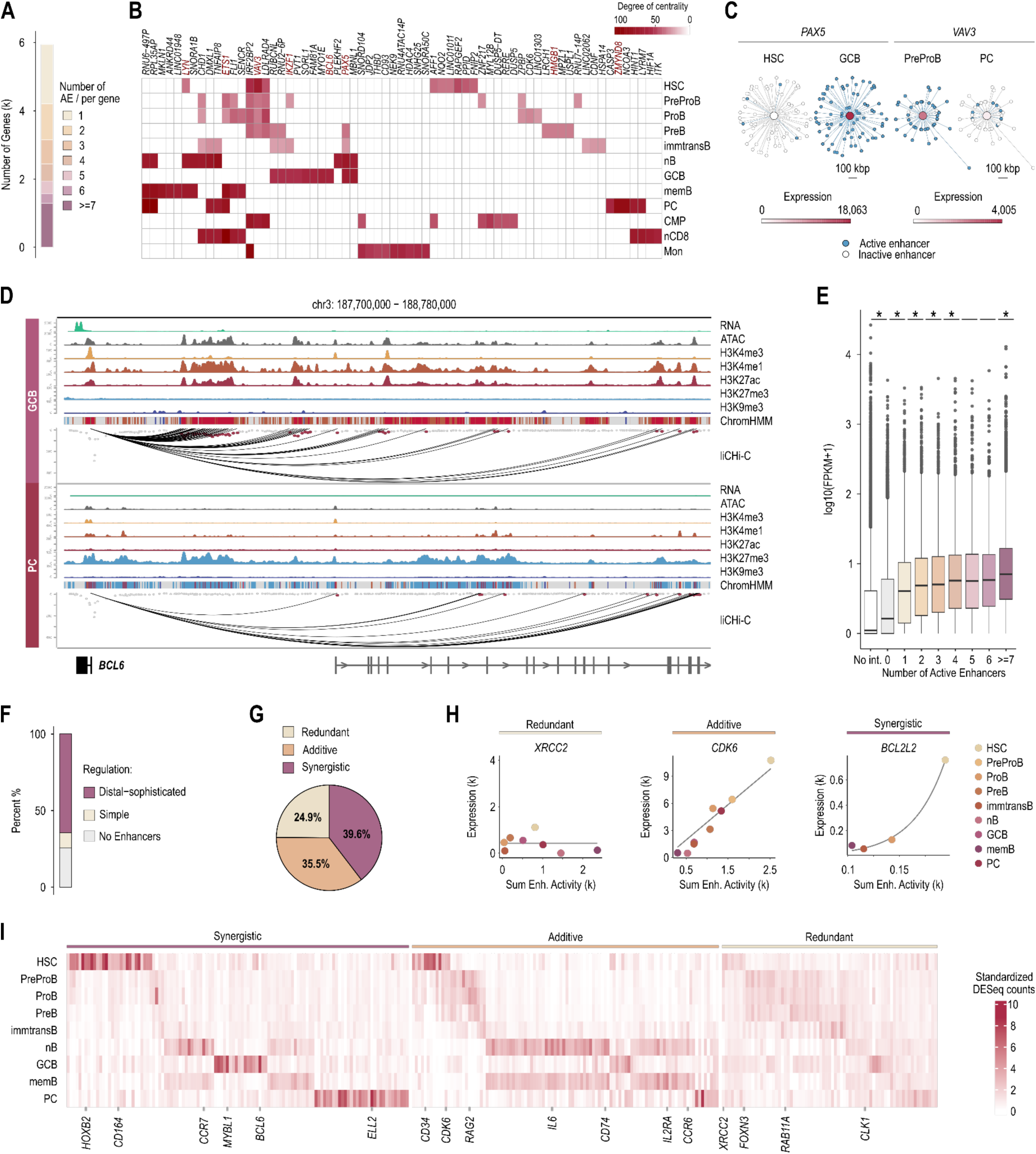
Additive and Synergistic Enhancer Interactions Coordinate the Regulation of B-Cell Fate and Function. **A.** Distribution of genes stratified by the number of promoter-linked active enhancers (AE), aggregated across all cell types. **B.** Heatmap showing the top 10 genes in each cell type based on degree centrality (number of direct connections). Columns are organized according to similarity in degree centrality between genes. Color intensity corresponds to absolute degree values (white = low, dark red = high). **C.** Gene-centric 3D enhancer networks of *PAX5 (left)* in hematopoietic stem cells (HSC) and germinal center B-cells (GCB); and of *VAV3 (right)* in PreProB cells and plasma cells (PC). Networks show AE in either of the two displayed cell types, arranged radially according to their genomic distance from the target gene (1 unit = 500 kbp) and connected by liCHi-C–detected interactions (grey edges). Enhancers are colored by activity in the indicated cell type (active = blue; inactive = white). Central node color represents *PAX5* and *VAV3* DESeq2-normalized expression levels in each cell type**. D.** *BCL6* regulatory landscape in germinal center B cells (GCB, top), and plasma cells (PC, bottom) according to RNA, open chromatin, histone modifications, ChromHMM states and liCHi-C data. **E.** Gene expression levels (FPKM) of genes stratified by the number of promoter-linked active enhancers, aggregated across all cell types. Asterisks indicate statistically significant differences (Wilcoxon test, p < 0.05) between adjacent groups. **F.** Distribution of protein-coding genes according to active enhancer regulation. Genes are classified as having no associated active enhancers (AEs) (grey), a single AE (*simple;* yellow), or multiple AEs (*distal-sophisticated*; purple). **G.** Pie chart showing the proportion of modeled genes classified as Redundant, Additive or Synergistic based on the best fitted model. **H.** Representative examples of Redundant (*XRCC2*), Additive (*CDK6*), and Synergistic (*BCL2L2*) genes. DESeq2-normalized RNA-seq expression is modeled as a function of the summed H3K27ac signal (Enhancer activity) across all interacting AEs for each gene and cell type. **I.** Heatmap of DESeq2-normalized expression levels for genes classified as Synergistic, Additive, or Redundant across all profiled cell populations. Expression values are standardized per gene and scaled across cell types by the gene’s mean expression. Representative genes from each subset are shown at the bottom.

In addition, genome-wide analyses showed a positive yet incomplete correlation between the number of active enhancers associated with a gene and either its expression level or promoter activity (Fig. 4E; Extended Data Fig. 11D), suggesting that enhancers may cooperate through diverse regulatory modalities to regulate B-cell fate and function. To gain deeper insight, we characterized the temporal functional cooperation of active enhancers, within their native genomic context and across the genome, to control gene transcription across B-cell differentiation. Our analysis revealed that approximately 25.6% of expressed protein-coding genes (FPKM > 5) are not associated with active enhancers and appear to be regulated exclusively through promoter activity, whereas 9.7% are controlled by a single active enhancer showing a simpler regulation (Fig. 4F). Functional enrichment analysis showed that genes governed by these simpler regulatory architectures are primarily involved in metabolic functions and rRNA-related processes, respectively (Extended Data Fig. 11E). In contrast, 65% of expressed protein-coding genes were linked to active enhancer networks, indicating that most genes are potentially co-regulated by multiple active enhancers acting simultaneously (Fig. 4F). Focusing on the last category, we modeled mRNA expression levels as a function of the cumulative H3K27ac signal across all linked active enhancers during B-cell differentiation and determined the best-fitting model using Bayesian Information Criterion (BIC) score (Extended Data Fig. 11F). 24.9% of genes that we were able to model were predicted to be regulated by redundant active enhancers, meaning that the gain of enhancer activity does not affect the expression of the target gene. Genes regulated by redundant active enhancers, exemplified by *XRCC2*, *RAB11A* and *FOXN3*, were primarily involved in cell division (Fig. 4G-H; Extended Data Fig. 11E,G). In addition, 35.5% of genes were regulated by additive active enhancers, meaning that each enhancer contributes proportionally to gene expression. Remarkably, genes regulated by additive active enhancers, exemplified by *CDK6*, *LEF1* and *IL6*, were enriched in lymphocyte processes, including cell differentiation, activation, and proliferation (Fig.4G-H; Extended Data Fig. 11E,G). Finally, 39.6% of modeled genes were regulated by synergistic active enhancers. This indicates that the combined effect of network enhancers on gene expression is substantially greater than the sum of their individual contributions, reflecting strong cooperative and hierarchical interactions among active enhancers. Interestingly, this set of genes includes both well-established regulators of B-cell biology—such as *BCL2L2 and ELL2*—and potential novel regulators identified in our study, including *MYBL1* (Fig. 4G-H; Extended Data Fig. 11G). Genes regulated by synergistic and additive active enhancers are more cell type–specific than those controlled by redundant active enhancers. Additive active enhancers tend to regulate genes expressed across multiple B-cell populations, particularly those active during early or late stages of differentiation, potentially contributing to B-cell fate acquisition. In contrast, genes regulated by synergistic active enhancers are typically highly expressed in a single B-cell population, likely reflecting their role in controlling population-specific functions or distinct steps of B-cell fate, which may explain why no single biological process showed significant enrichment (Fig. 4I; Extended Data Fig. 11H).

Collectively, these findings suggest that B-cell identity depends on more sophisticated regulatory input than common cellular processes, provided by large networks composed of active enhancers potentially acting additively and synergistically, enabling finely tuned transcriptional control of cell fate.

Until now we have presented the 3D epigenomic roadmap of human B-cell differentiation, which has enabled us to uncover the mechanisms through which active enhancers cooperate to drive human B-cell differentiation *in vivo* and to identify genes that represent candidate novel B-cell regulators. We next leveraged this dataset to demonstrate its potential for clinical translation, providing new insights that improve our understanding of both adaptive immunity—which enables rapid responses to repeated pathogenic challenges—and oncogenesis.

### Chromatin functions as a “3D memory machine”, enabling rapid B-cell recall responses to pathogenic challenges

Memory B cells provide long-term protection by rapidly differentiating into antibody-secreting cells upon re-exposure to their cognate antigen. This process, known as the immune recall response, forms the foundation of adaptive humoral memory^26^. Similarly to memory B cells, naive B cells are also able to differentiate into antibody-secreting cells upon encountering their cognate antigen for the first time (Fig. 1A). However, memory B cells respond more rapidly and efficiently than naive B cells, although the mechanisms underlying this faster recall response remain incompletely understood.

To address this gap, we first examined the possibility of a one-dimensional (1D) epigenetic priming mechanism at enhancers, which could predispose genes involved in proliferation, antibody diversification, and overall B-cell maturation to be more rapidly overexpressed upon secondary antigen recognition by memory B cells compared to their initial activation in naive B cells. To this end, we focused on protein-coding genes that are differentially upregulated in germinal center B cells relative to both naive B and memory B cells—but not differentially expressed between the latter—and designated these as “antigen-triggered B-cell genes” (Fig. 5A; Supplementary Table 4). Then we focused on those active enhancers that drive their high expression in germinal center B cells and studied the epigenetic state of those regions in naive B and memory B cells without considering the 3D chromatin organization. Overall, we observed a global conservation of chromatin states between naive B and memory B cells, with no gain of primed or poised states in the latter, but a slight increase in chromatin accessibility in memory B cells as previously reported in mice^27^ (Fig. 5B-C).

**Figure 5.**
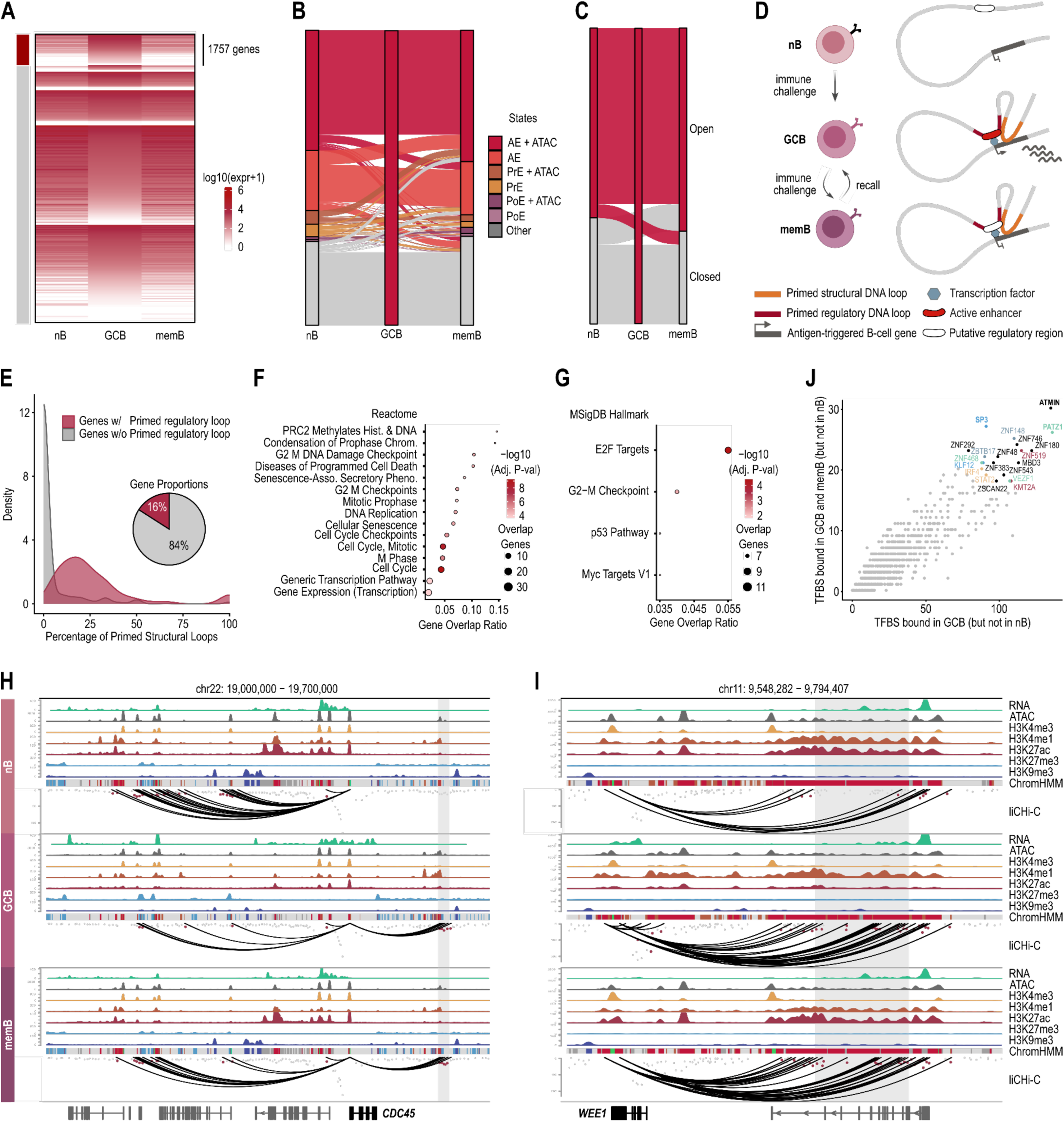
Primed Chromatin Loops Support Rapid B-Cell Recall Responses to previously encountered pathogens. **A.** Heatmap showing gene expression across naïve B cells (nB), germinal center B cells (GCB), and memory B cells (memB). Genes overexpressed in GCB cells (log2 fold change > 2 relative to both nB and memB) are highlighted by the red bar and were selected for downstream analyses. Heatmap colors represent log10-transformed expression values (log10[FPKM + 1]). **B.** Alluvial plot illustrating transitions in chromatin state annotations of active enhancers defined in GCB cells when tracked across nB and memB cells. States include active enhancers (AE), primed enhancers (PrE), poised enhancers (PoE), and other chromatin categories, with or without chromatin accessibility. **C.** Alluvial plot showing transitions in chromatin accessibility (open versus closed) of GCB-defined regulatory regions across nB, GCB, and memB cells. **D.** Graphical representation of both mechanisms by which the ‘3D memory’ is maintained in memory B cells: i) primed regulatory loops; and ii) primed structural loops. **E.** Left: Density distribution of the percentage of primed structural DNA loops per gene, stratified by the presence or absence of primed regulatory loops. Right: Pie chart showing the proportion of genes with and without primed regulatory DNA loops. **F.** Pathway enrichment analysis of genes associated with primed regulatory DNA loops using the Reactome database. Dot size represents the number of overlapping genes, color indicates −log10 adjusted P value, and the x-axis shows the gene overlap ratio. **G.** Pathway enrichment analysis of genes associated with primed regulatory DNA loops using the MSigDB Hallmark gene set collection, displayed as in (F). **H.** *CDC45* regulatory landscape in naïve B cells (nB, top), germinal center B cells (GCB, middle), and memory B cells (memB, bottom) according to RNA, open chromatin, histone modifications, ChromHMM states and liCHi-C data. **I.** *WEE1* regulatory landscape in naïve B cells (nB, top), germinal center B cells (GCB, middle), and memory B cells (memB, bottom) according to RNA, open chromatin, histone modifications, ChromHMM states and liCHi-C data. **J.** Predicted transcription factor binding sites (TFBSs) within GCB-specific active enhancers connected to antigen-triggered B-cell gene promoters through primed regulatory DNA loops. The x-axis represents TFBSs bound in GCB cells but not in nB cells, while the y-axis represents the subset of those TFBSs also bound in memB cells. Transcription factors are colored by motif cluster, with clusters defined based on motif similarity.

After excluding a 1D epigenetic priming as a major mechanism driving fast immune recall response, we investigated the possibility of a 3D epigenetic priming mechanism. To this end, we focused on the “antigen-triggered B-cell genes”, we identified the chromatin interactions linking active enhancers and promoters of these genes in germinal center B cells and examined loop dynamics in naive B and memory B cells independently of the 1D epigenetic landscape. We found that 16% of “antigen-triggered B-cell genes” have at least one “primed regulatory DNA loop”—defined as a loop connecting an active enhancer to an “antigen-triggered B-cell gene” promoter in germinal center B cells, conserved in memory B cells but not in naive B cells (Fig. 5D-E; Extended Data Fig 12A-B; Supplementary Table 4).

To evaluate the statistical robustness of this enrichment, we generated randomized gene sets and recalculated the proportions across thousands of permutations. The fraction of genes linked to “primed regulatory DNA loops” was significantly higher than expected by chance (p < 0.05), supporting a role for 3D epigenetic priming as a basis for B-cell immune recall (Extended Data Fig. 12C). In addition, genes with at least one “primed regulatory DNA loop” tend to preserve their overall structural landscape in memory B cells at a higher frequency than genes that only possess unprimed loops (Fig. 5E). Indeed, 41% of “antigen-triggered B-cell genes” have at least a “primed structural DNA loop”, a similar configuration to primed regulatory DNA loops but lacking an active enhancer in germinal center B cells. Primed regulatory DNA loops primarily regulate genes involved in cell proliferation, including MYC target genes, providing a mechanism to explain why memory B cells can initiate proliferation more rapidly than naive B cells upon antigen exposure (Fig. 5F-G). Examples of genes regulated by primed DNA loops include *CDC45*, which promotes DNA replication and S-phase progression; *WEE1*, a key enforcer of the G2/M checkpoint; and *MYBL1*, a cell cycle–regulating transcription factor that we identify as a novel regulator of germinal center B cells. Additional examples include *PCDHGA10*, *LHFPL2*, and *PAG1* among many others (Fig. 5H-I; Extended Data Fig. 12D-F).

Finally, we investigated the molecular basis of “primed regulatory DNA loops” by identifying TFs potentially contributing to their establishment and maintenance. We reasoned that TFs structurally regulating these loops should bind active enhancers engaged in such loops in germinal center B cells and remain bound in memory B cells but not naive B cells. However, TFs primarily involved in regulation of these genes—but not implicated in loop priming—should be predominantly bound to their active enhancers in germinal center B cells, where these genes are expressed, but not in naive or memory B cells. To test these hypotheses, we performed a footprinting analysis, which nominated several transcription factors as candidates, including PATZ1, ATMIN, and SP3. Notably, PATZ1 has recently been identified as a regulator of genome organization in mice, with roles in skeletal patterning and in corticospinal neurons during post-injury repair^28,29^, reinforcing its potential structural role in the establishment and maintenance of primed regulatory loops (Fig. 5J).

Although a functional demonstration—hindered by the inability to genetically perturb these transcription factors in humans during the differentiation of naive B cells into germinal center B cells and subsequently into memory B cells—is necessary to fully establish their structural role in loop formation and maintenance, this analysis demonstrates that chromatin acts as a “3D memory machine,” facilitating rapid B-cell recall responses to pathogenic challenges.

### Cancer Cells Retain Enhancer Signatures of Their Cell of Origin While Silencing Distal Regulation of Tumor Suppressor Programs

Given the well-established role of epigenetic deregulation in oncogenesis, we leveraged our 3D epigenomic roadmap of human B-cell differentiation to uncover new insights into tumor development. To achieve this, we utilized the International Human Epigenome Consortium (IHEC) EpiAtlas^30^—a multi-institutional reprocessing initiative that applies a standardized analysis pipeline to generate a uniform, high-quality, gold-standard collection of epigenomic maps. Specifically, we analyzed epigenetic data from seven B-cell malignancies, including leukemias, lymphomas, and myeloma, each originating from distinct B-cell populations at different stages of differentiation, as defined by their cell of origin (Fig. 6A). Based on the active enhancer set defined across human B-cell differentiation, we computed enhancer activity scores—as a proxy for enhancer activity—for each active enhancer–patient pair and applied Uniform Manifold Approximation and Projection (UMAP) to assess epigenetic similarities among patients and B-cell malignancy subtypes. Patients clearly clustered according to their type of malignancy and whether the disease originated during early B-cell differentiation (as in B-ALL) or late B-cell differentiation (as in the other malignancies), indicating that the active enhancer landscape faithfully reflects the underlying disease biology—in other words, each B-cell malignancy possesses a distinct “epigenetic signature” (Fig. 6B).

**Figure 6.**
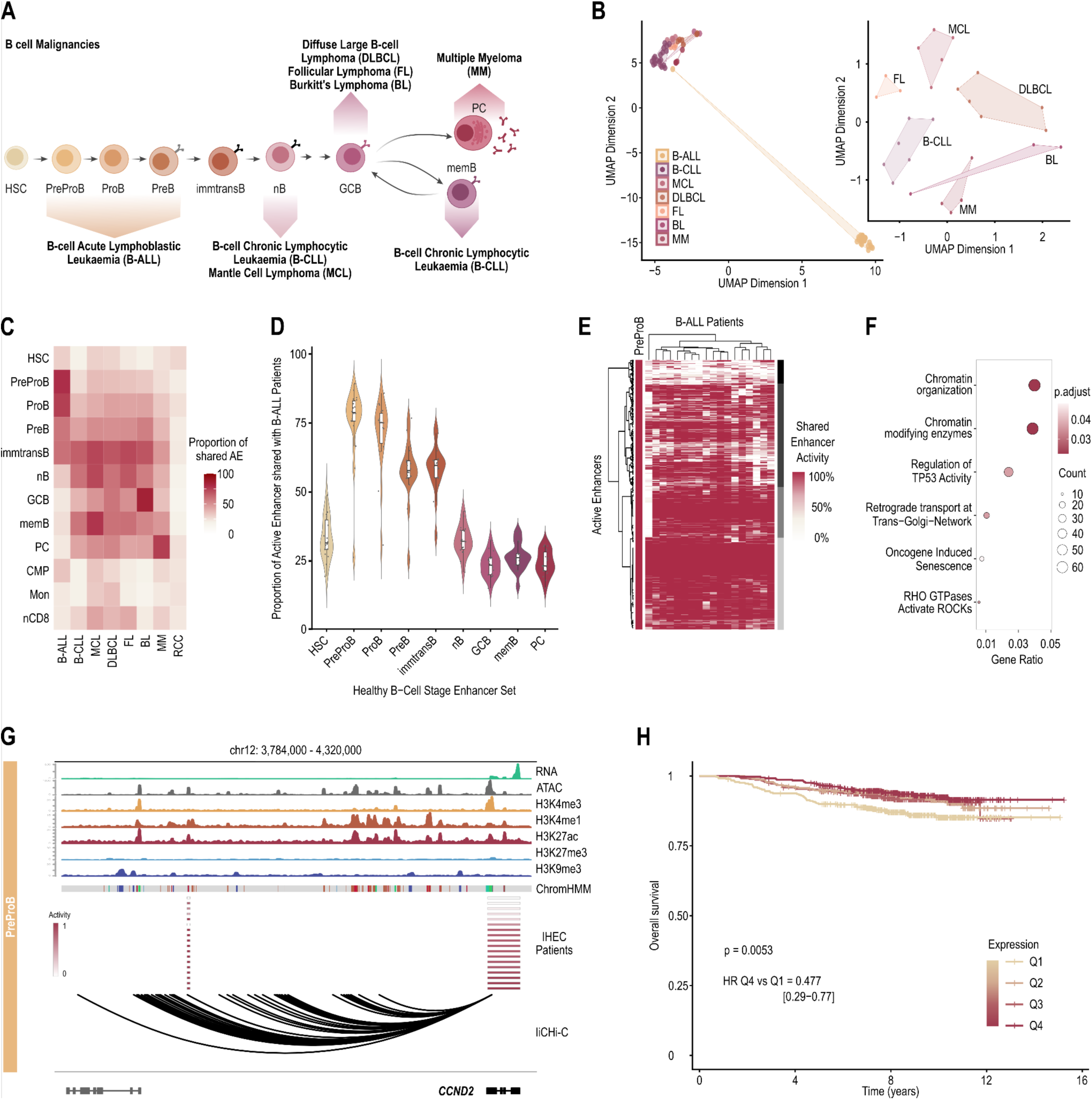
Malignant Cells Preserve Cell-of-Origin Enhancer Landscapes While Repressing Distal Regulation of Tumor Suppressor Pathways. **A.** Graphical representation of B-cell-derived malignancies along B-cell differentiation. **B.** UMAP of lymphoid malignancy patient samples based on enhancer activity scores profiles. Each dot represents a patient sample, colored by cancer type; convex hulls are shown for all groups to summarize each cancer type’s extent in UMAP space. Main: UMAP computed from cell-type–specific enhancer activity scores (“enhancer activity profile”); CLL downsampled (n = 18); seed = 42. Inset: UMAP computed from cell-type–specific enhancer activity scores; B-ALL excluded; CLL downsampled (n = 6); seed = 42. **C.** Heatmap of median cell-type–specific enhancer activity scores across lymphoid malignancies and renal cell carcinoma. Columns correspond to cancer groups ordered alphabetically, and rows correspond to cell-type–specific enhancer sets ordered according to B-cell developmental progression. Each tile shows the median enhancer activity score (percentage; 0-100) across patient samples within each cancer group for the corresponding enhancer set. Higher color intensity indicates a greater median fraction of enhancer activity attributable to that cell-type-specific enhancer set is present in the given cancer group. **D.** Enhancer activity profile for B-ALL across B-cell developmental enhancer sets. Violin plots show the distribution of enhancer activity proportions/scores across B-ALL patient samples for each healthy B-cell stage–specific active enhancer set (x-axis; ordered along B-cell developmental progression). Boxplots overlaid within each violin indicate the median and interquartile range, and jittered points represent individual patient samples. **E**. Heatmap of enhancer activity scores for PreProB–associated enhancers across B-ALL samples. Rows represent active enhancers that are annotated as present in the PreProB enhancer set (is_present_in_PreproB), and columns represent individual B-ALL patient samples (one column per sample). Each tile shows the per-sample enhancer activity score (range 0–1), displayed as a percentage (0–100%); higher values indicate greater activity for that enhancer in that sample. Values are shown without additional scaling, and both enhancers (rows) and samples (columns) are hierarchically clustered using Ward.D2. **F.** Reactome pathway enrichment analysis of genes linked to dysregulated active enhancers in B-ALL patients (black/dark grey clusters). Pathways shown are restricted to those exclusively enriched in this gene set following equivalent analyses of the other two clusters. **G.** *CCDN2* regulatory landscape in PreProB cells according to RNA, open chromatin, histone modifications, ChromHMM states and liCHi-C data. Enhancer (left) and gene (right) activity scores from B-ALL patient samples (IHEC) are shown for a representative active enhancer linked to and positively correlated with *CCND2* gene activity. **I.** Kaplan-Meier analysis of Overall Survival (OS) of MP2PRT-ALL cohort stratified by *CCND2* expression quartiles, displaying the Hazard Ratio (HR) between the highest (Q4) and lowest (Q1) quartiles.

To gain deeper insight into this “epigenetic signature,” we investigated whether it reflects the retention of the epigenomic memory of the cell of origin or the acquisition of a more immature epigenetic identity that drives malignancy. To this end, we compared the enhancer landscape of B-cell malignancies with that of each normal B-cell population, and as a proof-of-concept we used renal cell carcinoma (RCC) as a non-lymphoid tumor control dataset (Fig. 6C). In general terms, for each malignancy, the greatest similarity is observed with its corresponding cell of origin, sharing approximately 75% of the active enhancer landscape. For instance, B-ALL samples share on average 84% and 74% of active enhancers with PreProB and progenitor B cells, respectively—identified as their cells of origin—but less than 25% with unrelated populations such as germinal center B cells and plasma cells (Fig. 6D).

Notably, a subset of enhancers was inactivated in B-ALL samples compared to their healthy PreProB counterparts. Of these, 9.2% were constitutively inactivated in most patients (black), while 37.5% were inactivated only in a subset of B-ALL patients (dark grey) (Fig. 6E). To further explore this, we identified the candidate target genes of these inactivated enhancers using enhancer–gene associations derived from the cell of origin. Remarkably, these genes were enriched for pathways involved in the regulation of p53 activity and oncogene-induced senescence (Fig. 6F), suggesting that cancer cells may bypass intrinsic tumor-suppressive barriers by silencing their enhancers. In contrast, genes regulated by active enhancers conserved between cancer cells and their healthy counterparts were enriched for B-cell–associated pathways, including B-cell receptor and interleukin signaling, reinforcing the conservation of the enhancer landscape of the cell of origin (Extended Data Fig. 13B).

Finally, we leveraged epigenetic variation across B-ALL samples as a high-throughput, CRISPRi-like perturbation experiment to validate disease-associated gene findings and to provide genome-wide functional support for active enhancer–gene associations defined in the 3D epigenomic roadmap of human B-cell differentiation. To achieve this, we computed a gene activity score for each gene as a proxy for transcriptional output using epigenetic data. Focusing on enhancers inactivated in B-ALL patients relative to healthy counterpart, we correlated the activity scores of enhancers and their linked genes. This analysis validated 767 active enhancer–gene associations (FDR < 0.05) out of 11,347 tested. For instance, we identified *CCND2,* which encodes cyclin D2, a key regulator of cell-cycle progression, whose expression is associated with the overall survival of B-ALL patients (Fig. 6G-H). According to our data, the activity level of an enhancer located 340 kb upstream of the CCND2 promoter—previously identified as an active enhancer in PreProB cells—correlated with CCND2 transcriptional activity, providing orthogonal validation of this active enhancer–gene association.

Together, these findings show that cancer cells retain active enhancer signatures of their cell of origin while potentially silencing distal regulation of tumor suppressor programs, underscoring the power of our 3D B-cell epigenomic roadmap to pinpoint disease cells of origin—including those previously unknown—and identify cancer-associated genes, including novel ones.

### Collateral Damage from Gene Deletions Reveals a Novel Oncogenic Mechanism

Since not only epigenetic deregulation but also genetic alterations, such as deletions, are frequently associated with oncogenesis, we leveraged these alterations to extend the functional validation of active enhancer–gene pairs in an orthogonal manner. To achieve this, we leveraged matched genomic and transcriptomic data from B-cell malignancies—including B-ALL, diffuse large B-cell lymphoma (DLBCL), and multiple myeloma (MM)—to perform a CRISPR knockout–like perturbation analysis, enabling validation of the transcriptional regulatory roles of 213 deletions affecting active enhancers identified in the 3D epigenomic roadmap of human B-cell differentiation. (Fig. 7A). Among these, we identified several deregulated genes with oncogenic potential, including *COPS5*, *LPP*, *TCF4*, *TAP1*, *RPIA*, *VPREB1* and *SUV39H2*, a histone methyltransferase that mediates H3K9me3 and influences gene silencing and chromatin structure.

**Figure 7.**
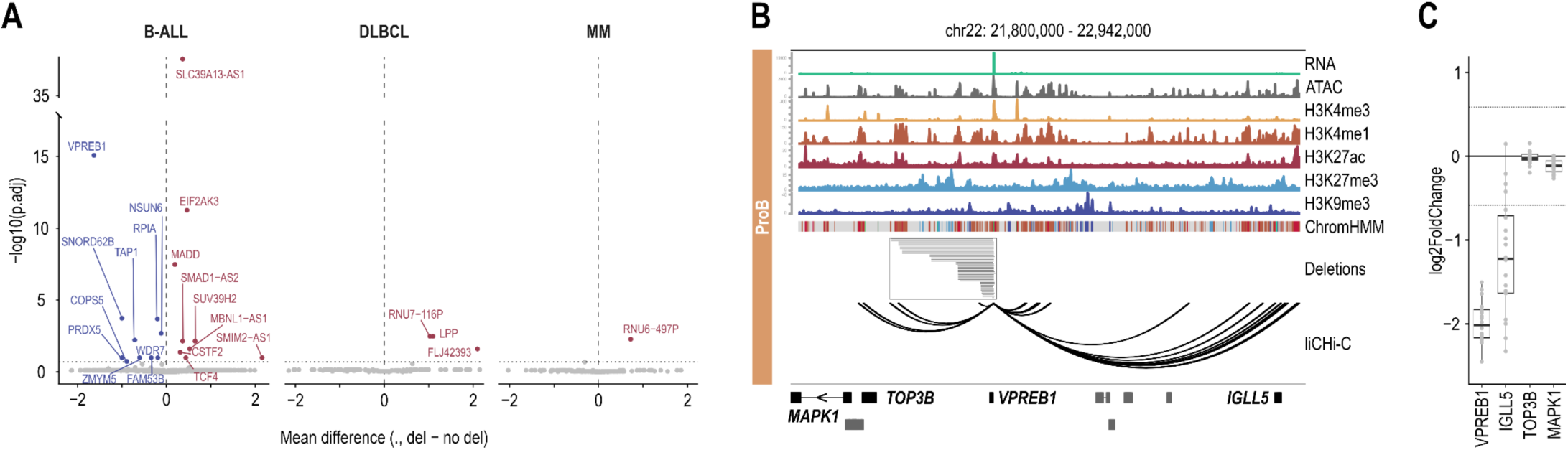
Intragenic Deletions Disrupt Distal Gene Regulation Through Enhancer Loss, Extending Beyond Simple Loss Of- Function Effects. **A.** Volcano plot summarizing the transcriptional impact on target genes whose promoter-linked active enhancers overlap genomic deletions. The x-axis represents the mean expression difference between samples carrying the deletion and samples without the deletion (Δ expression), and the y-axis indicates statistical significance. Each point corresponds to a gene; significant genes are highlighted. **B.** *VPREB1* regulatory landscape in progenitor B cells (ProB) according to RNA, open chromatin, histone modifications, ChromHMM states and liCHi-C data. The length of the deletions of *VPREB1* intragenic enhancers across the St. Jude B-ALL cohort are depicted in an independent track (grey shadow). **C.** Log2 fold change in expression of VPREB1 and distally interacting protein-coding genes (*IGLL5*, *TOP3B*, and *MAPK1*) between B-ALL samples carrying deletions affecting *VPREB1*-linked active enhancers and samples without such deletions in the St. Jude B-ALL cohort.

Since 81% of the enhancers we identified were located within gene bodies, yet only 33% of these intragenic enhancers were associated with their host gene (Fig. 1C), we hypothesize that gene deletions may contribute to oncogenesis not only through gene loss or the production of dominant-negative truncated proteins, but also through deregulation of distal target genes controlled by intragenic active enhancers—a phenomenon we term “collateral damage”. Indeed, 2.8% of deletions functionally validated affected intragenic active enhancers, supporting this previously overlooked mechanism of oncogenesis. To investigate this “collateral effect” in detail we focused on recurrent intragenic deletions associated with specific B-ALL subtypes. For example, deletions affecting the *VPREB1* gene — which encodes a component of the pre-B-cell receptor essential for early B-cell differentiation — occur in approximately 10% of B-ALL cases and are particularly frequent in pediatric B-ALL, affecting up to 33% of patients^31^. In progenitor B cells, multiple active enhancers—including some classified as super-enhancers—are located within intronic regions of *VPREB1* and, according to our data, regulate *IGLL5* rather than nearby genes such as *TOP3B* or *MAPK1*. Specifically, we identified focal *VPREB1* deletions, ranging from 3 to 232 kb in size, which resulted in the loss of intragenic active enhancers within *VPREB1* (Fig. 7B). In these cases, we observed more than a 1.5-fold decrease in the expression of *VPREB1* and *IGLL5*. *IGLL5,* encoding a surrogate light chain–like protein, supports progenitor to precursor B-cell transition, survival, and proliferation, and its dysregulation may contribute to the oncogenic transformation of progenitor B cells. In contrast, other genes located near the deleted enhancers but not regulated by them, such as *TOP3B* and *MAPK1*, showed no difference in expression between deleted and non-deleted cases (Fig. 7C).

In addition, we identified other notable examples of collateral damage mechanism from gene deletions, whereby intragenic deletions disrupt distal gene regulation through enhancer loss, extending beyond simple loss-of-function effects. One such gene is *ERG*, which is recurrently deleted in 3–7% of B-ALL cases and is highly enriched in patients with *DUX4* gene rearrangements ^32^. Historically, the primary consequence of these deletions has been associated with the generation of a truncated *ERG* protein with dominant-negative function. However, we found that in progenitor B cells, several active enhancers are located within intronic regions of *ERG*, which are frequently deleted (Extended Data Fig. 13B). According to our data, these active enhancers regulate *ETS2* and *PSMG1*. Supporting our proposed collateral damage mechanism, deletions of these enhancers were associated with more than a 1.5-fold decrease in expression of *ETS2*, *ERG*, and *PSMG1*. In contrast, no downregulation was observed for *KCNJ6*, which is not regulated by the deleted enhancers (Extended Data Fig. 13C). Interestingly, *ETS2* encodes a transcription factor highly expressed in B-cell progenitors that regulates proliferation, differentiation, apoptosis, and development, and can act as either an oncogene or a tumor suppressor depending on the context. In contrast, *PSMG1* is a proteasome chaperone with no direct established role in cancer.

Collectively, these results underscore the importance of looking beyond genomic hotspots—guided by 3D chromatin organization—not only to identify enhancer target genes, but also to understand the impact of genetic and epigenetic alterations in disease. Although this previously overlooked collateral mechanism of oncogenesis requires further investigation to assess its relevance across cancers, our findings underscore the value of integrating 3D B-cell epigenomic data across physiological differentiation stages to better elucidate the mechanisms driving malignant transformation and, ultimately, to inform the development of more effective therapeutic strategies.

## Discussion

Despite its central role in adaptive immunity and disease, the regulation of human B-cell differentiation remains poorly understood. This is mainly because most insights derive from mouse models, yet notable differences exist between the two species. For instance, mice display distinct phenotypic markers, differentiation stages, and signaling pathways, and are typically maintained in pathogen-free environments, limiting the physiological relevance of their immune responses^10^. Additionally, beyond genomic sequence divergence, species-specific regulatory programs are reflected in differences in enhancer activity, chromatin accessibility, and 3D genome organization. Therefore, while murine studies provide a valuable framework, especially for molecular interrogation, direct investigation of human B-cell differentiation is essential to reveal unique regulatory mechanisms that define human immune function and enable clinical translation of this knowledge to disease contexts. However, ethical constraints on tissue access, scarcity of early progenitors, and methodological challenges associated with limited cell numbers have hindered a comprehensive understanding of this process.

To address this limited understanding, several seminal studies have recently profiled open chromatin at single-cell resolution along human B-cell differentiation^33–37^. However, open chromatin primarily serves as a proxy for regulatory potential but does not accurately reflect the regulatory effect size of distal regulatory elements. Indeed, active enhancers, as well as poised or primed enhancers, and silencers all exhibit open chromatin. Moreover, these studies are limited in their ability to functionally link distal regulatory elements to their target genes, as they typically rely on correlations in chromatin accessibility between distal elements and nearby genes. A general limitation of such correlation-based approaches is that many predicted associations may represent indirect or secondary effects rather than *bona fide* cis-regulatory interactions. The use of 3D chromatin conformation data, however, facilitates this task. For instance, the Promoter Capture Hi-C^18,38^ and the Low Input Capture Hi-C (liCHi-C)^17,19^ methods that we developed, among others^39,40^, have been successfully used for the identification of disease-causative genes and pathways.

To overcome these knowledge gaps, we constructed a comprehensive 3D epigenomic roadmap of human B-cell differentiation by integrating histone modification and chromatin accessibility landscapes with 3D chromatin architecture mapped using liCHi-C, a low-input promoter capture Hi-C (PCHi-C) approach we recently developed, enabling the systematic identification of enhancer–gene associations ^17,19^. To our knowledge, this represents the most comprehensive and highest-resolution gene regulatory roadmap of a complete human cell differentiation series *in vivo* to date. Beyond providing detailed, novel insights into human B-cell differentiation and function in both health and disease, this resource also paves the foundation for uncovering general principles that govern human cell fate decisions. Importantly, this roadmap reveals that enhancers interact with their target genes over a median linear distance of 188 kb, with interactions extending beyond one megabase being relatively common. Moreover, most genes are likely regulated by multiple active enhancers. This highlights that approaches predicting enhancer–target gene associations based solely on linear proximity are inherently limited and may not fully capture the complexity of gene regulation.

Our analysis reveals that B-cell fate is governed by highly sophisticated gene regulatory mechanisms. In contrast to genes involved in general cellular processes, cell fate genes are regulated by large active enhancer networks that extend over roughly 1 Mb. These networks integrate multiple regulatory inputs, allowing precise and dynamic fine-tuning of gene expression over time. According to our findings, genes important for early or late stages of B-cell differentiation are regulated in a robust, additive manner to ensure developmental progression, whereas those defining fine-grained aspects of cell identity primarily rely on synergistic active enhancers, creating vulnerability points that are critical for precise B cell identity and may underlie disease when perturbed. Conversely, non–cell fate genes are controlled more simply by redundant active enhancers, or their regulation depends mainly on promoter activity or single active enhancers rather than distal enhancer input. Remarkably, although few, seminal studies have proposed gene-type–specific collaborative enhancer rules in *in vitro* systems across different species^41–43^; despite methodological variations, their findings broadly align with our observations in primary human samples along B-cell differentiation, further validating these principles. These regulatory patterns may mirror the evolutionary emergence of the genes themselves, with enhancer-independent or redundantly regulated architectures characterizing conserved cellular programs, and additive or synergistic active enhancer control underlying more recently evolved, specialized functions. Regardless of their potential evolutionary implications, this knowledge is essential for fully understanding the impact of both non-coding germline and somatic mutations, as well as non-coding epimutations, in genetically complex B cell–related diseases—such as leukemias, lymphomas, myelomas, autoimmune disorders, and immunodeficiencies—which are highly enriched in enhancer elements and may be driven by the cooperative behavior of perturbed enhancers.

In addition, we have discovered that chromatin acts as a “3D memory machine,” facilitating rapid B-cell recall responses to previously encountered pathogenic challenges. Consistent with a previous murine study showing that antiviral memory B cells possess numerous differentially accessible chromatin regions—mostly distal to promoters^27^—our findings confirm the conservation of this mechanism in humans and extend this concept to 3D. Importantly, a similar 3D epigenetic priming mechanism has been observed in CD4+ memory T helper 2 cells^44^, highlighting a conserved mechanism that enables rapid transcriptional immune responses. Here we do not propose a specific mechanism for the establishment and maintenance of primed regulatory DNA loops—which is energetically costly, as they must persist over extended periods, potentially spanning several years—as this lies beyond the scope of our study. However, we can speculate on two potential mechanisms: chromatin loop extrusion mediated by CTCF and cohesin and modulated by TFs or multivalent interactions between TFs and their cofactors that promote compartmentalization through condensate formation. In any case, TFs are the common denominator of both mechanisms, and our study identifies several candidates that may contribute. Notably, among these is PATZ1, a TF recently shown in mice to regulate genome architecture, whose loss leads to a reduction in DNA looping interactions ^28,29^.

Despite the value of our 3D epigenomic roadmap of human B-cell differentiation, several limitations remain. First, functional validation of active enhancers, as well as new factors regulating B-cell differentiation, is challenging due to difficulties in expanding and genetically manipulating primary B-cells, the lack of *in vitro* B-cell differentiation systems and interspecies differences. To partially address this limitation, we exploited cancer-associated alterations as a surrogate for large-scale functional validation in CRISPRi-like and CRISPR knockout–like frameworks. Second, B-cell differentiation is a continuous process characterized by transient states rather than discrete cell subtypes. Moreover, active enhancers and enhancer-promoter loops are often specific to subtypes and states. Because bulk samples inevitably comprise mixtures of these subtypes and states, our roadmap cannot fully capture the diversity, complexity, and continuous dynamics of enhancer regulation during human B-cell differentiation. Moving from bulk analyses to single-cell multi-omics approaches will be crucial for resolving state-specific enhancer–gene associations and characterizing developmental- and tissue-specific differences in B-cell differentiation, ultimately enhancing our mechanistic understanding of humoral immunity and B-cell–related diseases. Third, B-cell differentiation not only relies on the establishment of B-cell–specific transcriptional programs, but also on the silencing of alternative programs, such as myeloid or stem-cell programs. Here, we have only considered the role of active enhancers in B-cell differentiation, but an integrative study considering both enhancers and silencers will be fundamental to fully understand human B-cell differentiation in health and disease.

Despite its limitations, this pioneering 3D epigenomic roadmap of human B-cell differentiation represents an unprecedented resource for the field. First, it provides a powerful framework for identifying novel regulators of B-cell biology. For example, our study highlights MIXL1 and MYBL1 as a candidate regulator of germinal center B cells, whose functional role warrants further investigation beyond the scope of the present study. Notably, MYBL1 is also regulated by synergistic active enhancers forming a large active enhancer network and by primed DNA loops that facilitate rapid B-cell recall responses to pathogenic challenges. Second, although our analyses primarily focused on B-ALL, this resource holds broad potential to deepen understanding of other B-cell–related malignancies, autoimmune disorders, and immunodeficiencies. In particular, it enables exploration of how non-coding germline and somatic mutations, as well as epigenetic alterations, drive B-cell disease pathogenesis—ultimately paving the way for novel therapeutic approaches aimed at improving both life expectancy and quality.

## Materials & Methods

### Human Samples and Cell Isolation

A schematic diagram of the nine different cell states of the B-cell lineage isolated for this analysis can be found in Fig. 1A. Specifically, human HSC and B-cell progenitor populations were purified from fetal liver and fetal bone marrow of 12 to 22-week voluntary abortion donations, using immuno-magnetic columns and cell sorting. Briefly, single cell suspensions of fetal tissues were enriched in CD34+ and CD34-CD19+ fractions after a two-step immunomagnetic enrichment. First, CD34+ fraction was isolated using Miltenyi Biotec immuno-magnetic CD34+ enrichment kit (cat. 130-100-453) according to the manufacturer’s instructions. Second, the CD34- flow-through was used to isolate the CD34-CD19+ fraction using Miltenyi Biotec immuno-magnetic CD19+ enrichment kit (cat. 130-050-301) according to the manufacturer’s instructions. Then, CD34+ fraction was stained for cell sorting with 𝛼-CD34-PECy7 (BD Biosciences 348811), 𝛼-CD38-BB515 (BD Biosciences 564498), 𝛼-CD19-BV421 (BD Biosciences 562440), 𝛼-CD10-PerCPCy5 (BD Biosciences 563508), and 𝛼-CD33-PE (BD Biosciences 555450); and the CD19+ fraction was stained with 𝛼-CD19-BV421 (BD Biosciences 562440), 𝛼-IgM-PECy7 (Biolegend 314532), 𝛼-IgD-PE (Biolegend 307804), 𝛼-CD10-PerCPCy5 (BD Biosciences 563508), 𝛼-CD3-APC (Biolegend 317317), 𝛼-CD14-APC (Biolegend 398705), 𝛼-CD56-APC (Biolegend 392405). LIVE/DEAD aqua stain (ThermoFisher Scientific L34957) for viability using the manufacturer’s recommended concentrations (Suppl. Table 5). Finally, HSCs (CD34+, CD38-), CMPs (CD34+, CD38+, CD33+) and PreproB (CD34+, CD38+, CD19+, CD10-), progenitor B cells (CD34+, CD38+, CD19+, CD10+), precursor B-cells (CD19+, CD10+, IgM-, IgD-) and Immature-Transitional B (ImmtransB) (CD19+, CD10+, IgM+, IgD-/+) were sorted using a BD FACSAria Fusion flow cytometer with a 70 µm nozzle (Extended Data Fig. 1A).

Human naive B-cells, memory B-cells, naive CD8+ cells, and monocytes were isolated from buffy coat-peripheral blood mononuclear cells using immuno- magnetic enrichment kits following BLUEPRINT standard protocols. Particularly, naive B and naive CD8 isolation was performed using STEMCELL Technologies enrichment kits (cat. 19254 and 17968, respectively), while memory B cells and monocytes, were isolated with Miltenyi Biotec enrichment kit (cat. 130-093-546 and 130-091-765, respectively).

Human germinal center B-cells (CD19+, CD20+, CD38+) were isolated from routine pediatric tonsillectomies. Briefly, the tonsils were mechanically disaggregated and single live cells were separated from debris by Ficoll (Merck GE17-1440-03) gradient centrifugation. Then, single cell suspensions were enriched for B-cells using Miltenyi Biotec immuno-magnetic CD19+ enrichment kit (cat. 130-050-301) according to the manufacturer’s instructions. CD19+ pre-enriched cells were then stained for cell sorting with 𝛼-CD20-APCCy7 (Biolegend 302314), 𝛼-CD38-BB515 (BD Biosciences 564498) and DAPI (Merck D9542) for viability using the manufacturer’s recommended dilutions. Finally, GCBs were sorted using a BD FACSAria Fusion flow cytometer with a 100 µm nozzle (Extended Data Fig. 1A).

Plasma cells (PCs) were isolated from venous blood donations from healthy donors 7 days post-vaccination using the Miltenyi Biotec immuno-magnetic enrichment kit (cat. 130-093-628) according to the manufacturer’s instructions (Extended Data Fig. 1A). After cell isolation, sample purity was evaluated by FACS. Only those samples with a purity ≥90% of the population of interest were considered for subsequent analysis. Detailed information about magnetic microbeads and fluo-conjugated antibodies is available at Suppl. Table 5.

### Multi-omics Data Generation

#### CUT&RUN

For each histone post-translational modification (PTM), between 100,000 and 500,000 cells (depending on each cell type availability) were harvested in low-binding 1.5 mL tubes (Eppendorf 0030108051). Then, CUT&RUN protocol was performed as detailed in^45^ with the modifications detailed below. pAG-MNase was produced in-house by the Protein Science Facility (Karolinska Institute) and used at 700ng/ml. No *E.coli* spike-in was performed. The following antibodies were used to profile different histone PTMs: 𝛼-H3K27ac (Diagenode C15410196), 𝛼-H3K27me3 (Diagenode C15410195), 𝛼-H3K4me1 (Diagenode C15410194), 𝛼-H3K4me3 (Diagenode C15410003), 𝛼-H3K9me3 (Diagenode C15410193). Additionally, 𝛼-IgG (Diagenode C15410206) was used as a negative control (Suppl. Table 5). After chromatin digestion and release, DNA fragments were purified by phenol-chloroform extraction and used directly for library construction and amplification using the Kapa HyperPrep kit (Roche 07962363001) and Illumina’s TruSeq adapter system according to the manufacturer’s instructions. Adapters and PCR primers were synthesized by Integrated DNA Technologies; a list of the sequences for each adapter and primer can be found in Suppl. Table 5. CUT&RUN libraries were amplified by 10-11 PCR cycles, purified using 0.9x CleanNGS beads, and quantified and paired-end (PE) sequenced as detailed below.

#### ATAC-seq

Before starting, in-house produced Tn5 Protein (Science Facility, Karolinska Institute) was loaded at 4uM using Nextera XT Index Kit v2 (Illumina 15052163) compatible adaptors (Suppl. Table 5). First, equimolar amounts of Mosaic end_reverse were hybridized with Mosaic end_AdapterA or B using the following PRC program: 95°C 5min, ramping at -0.1°C/s to 25°, 25°C 5 min. Second, the following components (2µL 50uM MErev/ME-A, 2µL 50uM MErev/ME-B, 21µL Glycerol (100%, Sigma-Aldrich G5516), 2.79µL Tn5 (71.63 uM) and 22.21µL 2xLoading buffer (100mM Hepes (Sigma-Aldrich H4034), 200mM NaCl (Thermo Fisher Scientific Inc. AM9760G), 0.2mM EDTA (Thermo Fisher Scientific Inc. AM9260G), 0.2% Triton X-100 (Sigma-Aldrich T8787), 2mM DTT (Sigma-Aldrich D0632), 20% Glycerol (Sigma-Aldrich G5516) in water)) were mixed and incubated 1h at 25°C.

For each previously isolated population, between 10,000 and 100,000 cells (depending on each cell type availability) were harvested in low-binding 1.5 mL tubes. The ATAC-seq protocol was performed as detailed in^46^ with the modifications below. Tn5 was used at a final concentration of 0.328 uM and H_2_O topped up accordingly. After cell lysis, transposition and DNA purification, barcoded DNA-transposition fragments were used directly for PCR amplification using NEBNext Ultra II Q5 2× Master Mix (New England Biolabs M0544S) and Nextera XT Index Kit V2 (Illumina 15052163) following manufacturer’s instructions. ATAC-seq libraries were amplified by 10-14 PCR cycles, purified using 1x CleanNGS beads, and quantified and PE sequenced as detailed below.

#### RNA-seq

RNA was extracted from 70,000-200,000 cells using the RNeasy Mini RNA extraction kit (Qiagen 74104) according to the manufacturer’s instructions. Integrity and amount of the extracted RNA were assessed by automated electrophoresis using an RNA ScreenTape Analysis (Agilent 5067-5576) on an Agilent 2200 Tapestation. Only samples with 0.25-1 µg of total RNA and RINe value of 7 or higher were used for library construction. RNA samples were sent to Macrogen Inc. for mRNA library construction using SMARTer Ultra-Low RNAseq or TruSeq Stranded mRNA kits depending on available starting material. Sequencing was performed as detailed below.

#### liCHi-C

For each previously isolated population, between 500,000 cells and 1 million cells (depending on each cell type availability) were harvested in low-binding 1.5 mL tubes. The liCHi-C method was performed as detailed in^19^ with the modifications detailed below. Briefly, ligation products (pre-capture library) were amplified 9-10 cycles by PCR. Enrichment of promoter-containing ligation products was performed using SureSelectXT Target Enrichment System for the Illumina Platform (Agilent Technologies) as instructed by the manufacturer, and the library was amplified 7-8 cycles by PCR. Finally, the product was purified using 0.9x CleanNGS beads, and quantified and PE sequenced as detailed below.

#### Libraries Quality Control and Sequencing

Quantity and quality of the DNA libraries were assessed by automated electrophoresis using a high-sensitivity DNA ScreenTape Analysis (Agilent 5067-5584) on an Agilent 2200 Tapestation. The parameters used for discriminating between good and low-quality DNA libraries were based on the ones described in the literature for CUT&RUN^45^, ATAC-seq^46^ and liCHi-C^19^.

liCHi-C and CUT&RUN libraries were sequenced at Macrogen Inc. in South Korea using the HiSeq X 150+150 bp paired-end (PE) sequencing platform. liCHi-C libraries were sequenced to a minimum depth of 400M PE reads, while CUT&RUN samples were sequenced to a minimum of 30M PE reads. RNA-seq libraries were also sequenced at Macrogen Inc. using the NovaSeq6000 150+150 bp PE sequencing platform to a minimum of 40M PE reads. ATAC-seq libraries were sequenced at Novogene Co. in the United Kingdom using the Novaseq X Plus 150+150bp PE sequencing platform to a minimum of 250M PE reads. Final valid reads are available in Suppl. Table 1.

### Primary Data Processing Pipelines

#### CUT&RUN processing

PE reads were processed following the ENCODE standards^47^ with slight modifications. After trimming, reads were mapped using bowtie2 (2.3.2)^48^ to the reference genome (GRCh38.p13) with several adapted settings --very-sensitive-local --no-unal --no-mixed --no-discordant -k 2 --phred33 -I 10 -X 700 --dovetail. Filtering and deduplicating were performed using sambamba (0.7.0)^49^ and samtools (1.9)^50^ respectively. Bigwig files were generated using the function bamCoverage from deepTools (3.2.1)^51^.

Peak calling for the histone marks H3K27ac and H3K4me3 was performed using ChromTime (v1.0.0)^52^, which allows dynamic identification of chromatin state changes across temporal or pseudo-temporal trajectories. The analysis was conducted across eight developmental stages and along two differentiation trajectories terminating in memory B-cells and plasma cells. For stages shared between both trajectories, peak calls were intersected to increase robustness and reduce trajectory-specific variability. All analyses were performed using the GRCh38 reference genome. For each histone mark and stage, replicates were averaged, including both target (H3K27ac/H3K4me3) and corresponding IgG controls, prior to downstream processing. Sequence alignment files (BAM) were converted to BED format to meet ChromTime input requirements. ChromTime (v1.0.0) was run with default settings except for the following parameters, which were adjusted to increase stringency and improve reproducibility of dynamic peak detection: peak calling FDR (q-value) = 0.01, FDR seed (q-value-seed) = 0.01, and significance threshold for extending peak boundaries (p-value-extend) = 0.01.

#### ATAC-seq processing

PE reads were processed as described for CUT&RUN. The only extra step added was the offset correction of the paired read. Bigwig files were generated using the function bamCoverage from deepTools (3.2.1). Finally, peak calling was performed using HMMRATAC^53^ implemented in MACS3 v3.0.2, with default settings. Consensus peak files were obtained for each cell type by merging all peaks and keeping only those supported (partially or totally) by at least two replicates.

#### RNA-seq processing

PE reads were processed following the ENCODE standards^47^. First, reads were trimmed using Trim Galore (0.6.6) to remove any sequencing adapter. Then, reads were mapped to the reference genome (GRCh38.p13) using STAR (2.7.0f)^54^ with parameters recommended by ENCODE. Briefly, it filtered multi-mapped reads and tuned the allowed overhang and mismatches. Finally, read counts were quantified using featureCounts from subread (2.0.2)^55^, with the specific arguments -O (to handle reads falling into overlapping regions) and --countReadPairs (to count pairs of reads).

#### liCHi-C processing and interaction calling

PE reads were processed using HiCUP (0.8.2)^56^. First, the genome was computationally digested using the target sequence of the restriction enzyme. Then, the different steps of the HiCUP pipeline were applied to map the reads to the human genome (GRCh38.p13), filter out all the experimental artifacts and remove the duplicated reads and retain only the valid unique paired reads. To assess the capture efficiency, we filtered out those paired reads for which any end overlaps with a captured restriction fragment, retaining only the unique captured valid reads for further analysis. Library statistics for all samples are presented in Suppl. Table 1.

Interaction confidence scores were computed using the CHiCAGO R package (1.14.0)^57,58^. In summary, this pipeline implements a statistical model with two components (biological and technical background), together with normalization and multiple testing methods for capture Hi-C data. CHiCAGO analysis was performed in merged samples to increase the sensitivity, after assessing for reproducibility between biological replicates using principal component analysis, and hierarchical clustering. Significant interactions with a CHiCAGO score ≥5 were considered as high-confidence interactions. Interaction statistics for all samples are presented in Suppl. Table 1.

#### Clustering of promoter interactions

Interactions were clustered using k-means algorithm and for each cluster a specificity score was computed using the asinh-transformed CHiCAGO scores^18^. Clustering of cell types was performed using a hierarchical method with average linkage based on Euclidean distances, and principal component analysis was performed using the prcomp function in R. For interaction data handling we used HiCaptuRe Bioconductor R package^59^ which allows us to integrate epigenomic and transcriptomic data with liCHi-C or generate virtual 4C of specific genes.

#### Data Quality Control and Replicates selection

To ensure high-quality CUT&RUN and ATAC-seq datasets, library complexity, fragment size distribution, and signal-to-noise ratio were assessed. Library complexity was evaluated using three metrics recommended by the ENCODE consortium—Non-Redundant Fraction (NRF), PCR Bottlenecking Coefficient 1 (PBC1), and PCR Bottlenecking Coefficient 2 (PBC2)—computed with an in-house script; higher values indicate greater complexity and lower PCR bias. Fragment size distributions were calculated from mapped BAM files using samtools-based scripts, with ATAC-seq libraries expected to show peaks corresponding to nucleosome-free and nucleosomal fragments and CUT&RUN libraries primarily showing a mononucleosome peak (∼147 bp). Signal-to-noise ratio was estimated using the fingerprintPlot function from deepTools, summarizing the genome-wide read distribution via the area under the curve (AUC), where strong signal enrichment corresponds to lower AUC values. Libraries failing to meet the expected thresholds in any of these metrics were flagged for potential exclusion or interpreted with caution in downstream analyses.

#### Chromatin Segmentation and State Annotation

Chromatin states were identified using ChromHMM (v1.24)^60^ based on CUT&RUN profiles for five histone PTMs: H3K4me1, H3K27me3, and H3K9me3 (broad histone PTMs), and H3K4me3 and H3K27ac (narrow histone PTMs). For B-cell populations (HSC, PreproB, ProB, PreB, ImmtransB, nB, GCB, MemB, and PC), two types of inputs were generated depending on the histone PTM. Broad histone PTMs were binarized using the *BinarizeBam* function, whereas narrow histone PTMs were binarized using the *BinarizeBed* function from peak calls generated with ChromTime. For each cell type and chromosome, the resulting binary files for the five histone PTMs were merged into a single binarized matrix, with one column per histone PTMs. Control cell types (CMP, Mon, nCD8) were processed similarly, using the *BinarizeBam* function for all histone PTMs.

A 15-state ChromHMM model was trained with the *LearnModel* function using only the binarized datasets from B-cell types. This trained model produced genome-wide chromatin state segmentations for all B-cell populations. The resulting model was then applied to the control cell types to generate their corresponding segmentations using the *MakeSegmentation* and *MakeBrowserFiles* functions. The emission probability matrix produced by the trained model was subsequently annotated to assign functional chromatin states. Prior biological knowledge was used to reinterpret and relabel the 15 states into biologically meaningful chromatin state categories (Extended Data Fig. 2B). All cell-type–specific segmentation files were updated accordingly.

Finally, the annotated segmentation files were integrated with cell-type–specific ATAC-seq peaks obtained from HMMRATAC. A disjoin approach was applied to subdivide chromatin segments based on their overlap with ATAC peaks, enabling precise identification of accessible regions within each chromatin state.

#### Super-Enhancer Identification

Super-enhancers were identified for each cell type using the Ranking of Super Enhancers (ROSE) algorithm^61,62^ with default parameters. CUT&RUN H3K27ac and IgG BAM files were used as input and control files, respectively. As constituent enhancers, we used the active enhancers regions overlapping ATAC-seq peaks.

#### Assessment of Active Enhancers

Active enhancers were identified in each cell population using ChromHMM and ATAC-seq peaks, and target genes were subsequently assigned using liCHi-C. Since active enhancer regions differ in size and location across cell types, the H3K27ac signal was quantified using a genome-wide binarization approach with 600bp bins, approximating the median size of linked active enhancers. PCA was then performed on these active enhancers overlapping bins linked to at least one target gene, allowing comparison of enhancer activity patterns across cell populations.

#### Definition of Cell-Type- Specific and Shared Active Enhancers

Active enhancers were considered specific when they did not overlap with active enhancers detected in any other cell type. For developmental groups, active enhancers were classified as group-specific based on active enhancer overlapping across related cell populations. Early B-specific active enhancers were defined as those shared by at least two of the following cell types: PreProB, progenitor B, precursor B, and immature-transitional B cells. Late B-specific active enhancers were defined as those shared by at least two of naive B, memory B, germinal center B, and plasma cells. B-cell-specific active enhancers were defined as those shared in at least five B-cell-related cell types. For each subset of specific active enhancers, putative target genes were assigned using promoter–enhancer interactions derived from liCHi-C data.

#### Transcription factor footprinting

Transcription factor (TF) motifs were obtained from HOCOMOCO v13 (HUMAN)^63^. Only primary (subtype = 0) and high-quality (A–C) motifs were retained. Gene symbols were standardized according to UniProt and manually corrected where necessary. Motifs were further filtered by expression using DESeq2-normalized RNA-seq counts from B cell developmental populations, retaining only TFs with a median normalized count ≥ 5 in at least one cell type. The resulting expressed TF motif set was used for footprinting analysis the TOBIAS framework (v0.17.0)^64^ in three sequential steps: 1) bias correction of ATAC-seq reads using ATACorrect, 2) calculation of footprint scores from bias-corrected cut sites within accessible peaks with ScoreBigwig, and 3) identification of TF binding using BINDetect. Each step was run per cell type using corresponding ATAC-seq BAM files and peak sets.

#### Enrichment of transcription factor binding

The output from the TOBIAS footprinting analysis was used to detect enrichment of specific TFs in primed regulatory DNA loops and in cell type-specific enhancers. To detect TFs enriched in cell type-specific enhancers, TF binding was quantified for each cell type in cell type-specific versus non-cell type-specific enhancers. For each TF we calculated the percentage of specific and non-specific enhancers containing at least one bound site. A linear model was fitted relating TF binding frequency in specific enhancers to its binding frequency in non-specific enhancers. Positive residuals (observed minus expected specific binding) represent TFs bound more often than predicted to cell type-specific enhancers. TFs were ranked by residuals, and the top 10 per cell type were selected. TFs with a median normalized expression levels below 5 were removed. For some plots, TFs were grouped in clusters based on the grouping by motif similarity downloaded from HOCOMOCO v13.

To detect TFs enriched in primed regulatory DNA loops, we calculated occupancy of each TF in these loops (percentage of loops with at least one bound TFBS) and then checked what percentage of these bound TFBS were maintained in memory B cells. Only TF expressed in both cell types (germinal center B-cells and memory B-cells) were considered (minimum 1 normalized gene counts).

#### Active Enhancer Networks

Gene-centric 3D active enhancer networks were constructed using the R package igraph (v.2.1.4)^65^ and the cell-type specific interactions pairs between genes and active enhancers obtained from liCHi-C.

Networks were built for all cell types using non-directed graphs, considering genes as central nodes. Degree of centrality per gene was calculated using the function degree from igraph (v.2.1.4)^65^. Genes interacting with only a single active enhancer were excluded from the analysis as they provided limited information. The degree of centrality heatmap was created using the R package pheatmap (v1.0.13)^66^.

Active enhancer networks were visualized with custom radial plots. Edges lengths were plotted based on the normalized genomic distance between active enhancers and genes (1 unit = 500 Kbp). Enhancers shared between the compared cell types were drawn once.

#### Gene Expression-Active Enhancer Regulatory Modeling

Gene expression of distal-sophisticated genes (protein-coding genes interacting with more than one active enhancer) was modelled as a function of enhancer activity, following a framework adapted from^67^ and^68^. For each gene and each B-cell–related cell type, mean values across biological replicates of DESeq2-normalized RNA-seq expression were quantified, together with the mean summed H3K27ac signal (DESeq2-normalized signal) across all interacting active enhancers. For each gene, three alternative regulatory models were fitted to relate gene expression to enhancer activity.

Additive: RNAgene=i=1nH3K27ac Enhi +

Synergistic: RNAgene= expi=1nH3K27ac Enhi +

Redundant: RNAgene=

where 𝛼 represents the effect size of enhancer activity and 𝛽 the basal expression level. Protein-coding genes whose associated active enhancers were present in only one or two cell types were excluded, as this provided insufficient observations to reliably fit the regulatory models. Model selection was performed using the Bayesian Information Criterion (BIC). Relative BIC values (ΔBIC) were computed by comparing the Additive model against the Synergistic (*ΔBIC Add–Syn = BIC Additive − BIC Synergistic*) and Redundant models (*ΔBIC Add–Red = BIC Additive − BIC Redundant*). Genes with ΔBIC Add–Syn > 6 and ΔBIC Add–Red ≤ 0 were classified as Synergistic, whereas genes with ΔBIC Add–Red > 6 and ΔBIC Add–Syn ≤ 0 were classified as Redundant. Genes with negative ΔBIC values against both Synergistic and Redundant models were classified as Additive. A stringent ΔBIC threshold of 6 was applied to ensure robust model selection, prioritizing clear regulatory behaviors and minimizing ambiguous or potentially misleading classifications. Genes failing to meet these criteria or showing conflicting support across model comparisons were considered ambiguous and excluded from downstream analyses.

#### Analysis of Primed Regulatory DNA Loops

To assess the significance of the observed proportion of genes with primed regulatory DNA loops, a bootstrap analysis was performed. Genes that were not overexpressed in germinal center B-cells were used as the control set. For each bootstrap iteration, a random sample of control genes equal in size to the number of overexpressed genes captured by liCHi-C was selected. The proportion of genes with primed regulatory loops was calculated using the same criteria applied to the overexpressed gene set. This procedure was repeated for 2,000 iterations to generate a null distribution of proportions. An empirical p-value was then estimated as *p=((number of bootstrap proportions>observed proportion)+1*) / (number of iterations+1)

#### IHEC EpiAtlas analysis

##### Chromatin state data and sample selection

Chromatin state profiles for human epigenomes were retrieved from the International Human Epigenome Consortium (IHEC). ChromHMM emission matrices were downloaded per chromosome and processed from the standardized ChromHMM matrix directory for each epigenome. Sample metadata, including harmonized disease labels and lineage assignments, were obtained from the IHEC Metadata Harmonization File (v1.3). Only lymphoid-lineage cancer samples were retained, excluding all T-cell–derived samples. The final dataset comprised 158 B-cell malignancy samples, including 18 B-acute lymphoblastic leukemia (B-ALL), 3 Burkitt lymphoma (BL), 6 diffuse large B-cell lymphoma (DLBCL), 3 follicular lymphoma (FL), 5 mantle cell lymphoma (MCL), 119 chronic lymphocytic leukemia (CLL), and 4 multiple myelomas (MM).

##### Definition of cell-type enhancers and cell-type–specific enhancers

Enhancer annotations were obtained from the specific active enhancer catalog. Two complementary enhancer sets were defined for each B-cell developmental stage:

- *All cell-type enhancers:* all enhancer loci annotated as active in a given B-cell stage, regardless of whether they are also active in other hematopoietic or non-hematopoietic cell types.
- *Cell-type–specific enhancers:* the subset of enhancers annotated as uniquely active in that B-cell stage and absent from all other profiled cell types.

Enhancer coordinates were expanded into 200-bp genomic bins. Each bin was assigned to its corresponding B-cell stage and enhancer category (all vs. cell-type–specific) based on the specific active enhancer specificity annotations. This produced a non-overlapping bin-level enhancer map spanning early B-cell progenitors (e.g., HSC/early progenitor, progenitor B-cells) and late stages (e.g., germinal center B cells, plasma cells).

##### Extraction of ChromHMM states for enhancer bins

For each chromosome, ChromHMM segmentation matrices were loaded and aligned to the enhancer bin map. For every enhancer-associated bin, the ChromHMM state was directly extracted from the corresponding genome-wide segmentation. ChromHMM enhancer-related states were defined following the original model (states 3–4 and 7–11), representing strong and weak enhancer activity.

##### Enhancer activity score (enhancer state proportion)

Enhancer activity for each sample and each B-cell stage was quantified as the proportion of enhancer bins annotated with an enhancer ChromHMM state:

Enhancer Activity Score = # enhancer bins in enhancer states# enhancer bins Scores were computed at two complementary resolutions:

- *Sample-level score*: all bins from a given enhancer set were concatenated for that sample. Depending on the analysis, this included either:
  - all enhancers active in that B-cell stage, or
  - only the cell-type–specific subset.

This yielded a single activity score per sample per developmental stage.

- *Enhancer-level score*: the score was computed for individual enhancers, using only the bins belonging to that enhancer in that sample.

##### Gene activity score

As a proxy for gene activity, an analogous score was computed using bins annotated with any active transcription- or enhancer-associated state (ChromHMM states 1–11). This yielded a per-sample Gene Activity Score reflecting overall chromatin activation across gene-associated regions.

#### Structural Variant and Enhancer Deletion Analysis

##### Genome-wide analysis of enhancer deletions impact on gene expression

The NIH Database of Genotypes and Phenotypes (dbGaP) was used to access whole genome sequencing (WGS) datasets for deletions analysis. Specifically, we included 2373 participants across 3 B-cell malignancies: 1468 B-cell Acute Lymphoblastic Leukemia (B-ALL) cases from The Molecular Profiling to Predict Response to Treatment project (MP2PRT, dbGaP accession: phs001965.v1.p1), 695 Multiple Myeloma (MM) from The Multiple Myeloma Research Foundation Relating Clinical Outcomes in MM to Personal Assessment of Genetic Profile trial (MMRF-COMMPASS, dbGaP accession: phs000748.v7.p4), 74 Diffuse Large B-cell Lymphoma (DLBCL) from The Cancer Genome Atlas (TCGA-DLBC, n= 42, dbGaP accession: phs000178.v11.p8) and Cancer Genome Characterization Initiative HIV+ Tumor Molecular Characterization Project (CGCI-HTMCP-DLBCL, n=32, dbGaP accession: phs000235.v21.p6). For somatic deletions analysis we included only those cases for which structural variants (SVs) had been called using Manta^69^ and corresponding RNA-seq data from the same patient were available. Patients with more than 1,715 structural variants passing all quality filters were excluded as hypermutated, corresponding to samples with more than three standard deviations above the mean log₁₀-transformed count of high-confidence variants.

Somatic deletions were intersected with defined active enhancer regions to identify patients with disrupted regulatory elements. Patients were stratified into “Affected” (deletion carriers) and “Unaffected” (wild-type) groups based on the presence of a somatic deletion within any enhancer linked to a given tested gene in cell-type of origin (PreProB, ProB, PreB, immtransB for B-ALL, GCB for DLBCL, PC for MM). The impact on the expression of linked target genes was assessed by comparing variance-stabilized (VST) RNA-seq counts between patients harboring deletions and unaffected controls using Wilcoxon test. The effect size was calculated as the difference in mean expression (Δ) between the deleted and non-deleted groups. P-values were adjusted for multiple testing using the Benjamini-Hochberg (BH) false discovery rate (FDR) method.

##### Analysis of *ERG/VPREB1* intragenic enhancer deletions

SNP array DNA copy number (CN) and RNA-Seq HTSeq counts were extracted from a cohort of 1,988 B-ALL patients^70^, available via the St. Jude Cloud at https://viz.stjude.cloud/st-jude-childrens-research-hospital/collection/pax5-driven-subtypes-of-b-progenitor-acute-lymphoblastic-leukemia~1. Patients with log2 CN ratios < -0.43 at *ERG* (n = 12) and *VREPB1* (n = 39) progenitor B-cells active enhancer region, in the absence of further significant local CN alterations, were cross-referenced with RNA-Seq HTSeq count data.

20 random permutations of RNA-Seq counts from an equivalent number of patients without *ERG* (n = 12) or *VPREB1* (n = 39) intragenic enhancer CN deletions, were extracted. DESeq2 (v1.48.2) was then used for differential gene expression analysis between *ERG* and *VPREB1 intragenic e*nhancer deleted patients and each of the 20 random samples of non-CN–loss patients. DESeq2 differential expression results were then filtered for genes shown to interact with or located near *ERG* or *VPREB1* intragenic enhancers in progenitor B-cells, as determined by liCHi-C. Fisher’s test was implemented using the poolr R package (v1.2.0) to determine overall statistical significance across DESeq2 permutations for each gene, and effect size cut-off of 1.5 linear fold-change (0.585 log2 fold-change) in either direction was employed.

#### Survival Analysis

RNA-seq counts of MP2PRT Cohort data^71^ (n=1479) were normalized using the DESeq2 package. Principal Component Analysis (PCA) was performed on variance-stabilized transformed (VST) counts to assess sample clustering and ensure the absence of outliers. For survival analysis, patients were stratified into quartiles based on normalized gene expression levels of candidate CCND2 gene. Kaplan-Meier curves were generated to estimate Overall Survival, with differences assessed via the log-rank test. Hazard Ratios (HR) comparing the highest (Q4) versus lowest (Q1) expression quartiles were calculated using univariate Cox proportional hazards regression.

#### Gene Ontology and Pathways Enrichment Analysis

Gene Ontology (GO) and pathway enrichment analyses were performed on gene sets derived from enhancer–gene interaction analyses or differential expression comparisons using GO Biological Process (BP) terms or curated pathway databases. Statistical significance was assessed using multiple-testing correction, with an adjusted p-value (FDR) < 0.05, and enriched terms were visualized using dot plots. For genes interacting with specific active enhancers, significant GO terms were manually grouped into functional categories and clustered based on gene overlap. Pathway enrichment for the 3D-primed machinery analysis was performed using Reactome and MSigDB via Enrichr on genes strongly upregulated in germinal center B cells (log2FC > 2 relative to both naive B cells and memory B), without specifying a background gene universe. Reactome pathway enrichment was also applied to genes interacting with dysregulated active enhancers, retaining pathways that were significant and unique relative to those obtained from conserved enhancer interactions. All protein-coding genes were used as the background universe for specific active enhancer and dysregulated enhancer analyses, whereas only the expressed ones (>5 FPKM) were used for enhancer regulatory mechanism analyses.

#### Epigenome Browser Data Visualization

Gene landscape profiles were generated using an in-house script based on the Gviz R package^72^. For each example gene, all promoter interactions and CHiCAGO interaction outputs were displayed. To enable consistent comparison across cell types, scale limits for each omics dataset were set equally across cell types. RNA-seq, ATAC-seq, and CUT&RUN profiles for all histone marks were visualized from BigWig files, applying a smoothing method to improve clarity. For gene tracks, all transcripts of a given gene were collapsed into a single fragment spanning the entire gene region, aggregating all transcript coordinates.

## Data Availability

The data that support this study are available from the corresponding author upon reasonable request. All raw omics datasets generated in this study have been deposited in EGA under the accession number **(pending)**. These datasets will be shared with controlled access in accordance with the ethical consent signed by the volunteers and it is limited to not-for-profit organizations after providing documentation of local IRB/ERB approval. Data Access Committee, led by Biola M. Javierre, will determine access permissions in a timeframe of fifteen workdays. The processed files from all the omics datasets have been deposited in Gene Expression Omibus under the accession number **(pending)**.

## Code Availability

All analysis done in the present article are accessible in the GitHub repository: https://github.com/JavierreLab/B-Cell_Roadmap

## Acknowledgments

We gratefully acknowledge the participation of all volunteers. We are indebted to the Blood and Tissue Bank of the Catalan Healthcare System; the Department of Otorhinolaryngology at Germans Trias i Pujol Hospital; and the Department of Hematology at Germans Trias i Pujol Hospital−especially Encarnación Santafé, Carmen Villena and Isabel Granados—for providing the non-fetal samples. The human embryonic and fetal material was provided by the Human Developmental Biology Resource (Project Reference 200640). We are also grateful to the Flow Core Units at the Institute for Health Science Research Germans Trias i Pujol–Josep Carreras Leukaemia Research Institute, Newcastle University, and the Scientific and Technological Center of the University of Barcelona. We also thank the members of the Javierre Group for their valuable discussions. Some illustrations were partially created using BioRender (https://biorender.com). This work was supported by the La Caixa Banking Foundation (LCF/PR/HR23/52430008), FEDER/Spanish Ministry of Science and Innovation (PID2021-125277OB-I00), the Wellcome Leap HOPE programme (21-275 RCA), the Deutsche José Carreras Leukämie Stiftung (DJCLS 18 R/2023), the 2024 Bilateral Grant from the European Hematology Association, and the 2024 Leonardo Grant for Scientific Research and Cultural Creation from the BBVA Foundation. LT-D is funded by the FPI Fellowship (PRE2019-088005), NB-A and LR are funded by the AGAUR FI fellowship (2025 FI-100864 and 2019 FI-B00017, respectively), PL-M is funded by the FPU fellowship (FPU20/03798 and EST24/00286), MR is funded by the CarrerasLeader-HORIZON-MSCA-2021-COFUND (GA 101081347), GM is funded by CarrerasPathfinders-HORIZON-MSCA-2022-COFUND-Doctoral (GA 101126688), LA is funded by MICIU/AEI/10.13039/ 501100011033 and FSE+ (JDC2024-053347-I), TR-F is funded by AECC-Talent-HORIZON-MSCA-COFUND (TALEN246598ROBE), AP-R is funded by MICIU/AEI / JDC2022-049753-I), DR is funded by FEDER/Spanish Ministry of Science and Innovation (PID2023-148272OB-I00), and BMJ is funded by the Spanish Health Institute Carlos III (CP22/00127). The funder bodies were not involved in the study design, collection, analysis, interpretation of data, the writing of this article or the decision to submit it for publication.

## Institutional Review Board Statement

The study was conducted according to the guidelines of the Declaration of Helsinki and approved by the Institutional Review Board of the Clinical Research Ethics Committee of University Hospital Germans Trias i Pujol (REF.CEI: PI-23-174, PI-23-251).

## Author contributions

RdH-B, LT-D, LF-E, and BMJ designed the research in collaboration with LJR and DR. RdH-B, LT-D, P L-M, NB-A, JO, MR, GM, LA, JC, AA, NB, JF-B, AC, and SP developed computational frameworks and performed data analysis. LF-E, LR, BV-M, BU, AP-R, and TR-F conducted the experiments. DG and EA provided samples. JM-S and AV offered critical feedback. BMJ secured funding, led the project and wrote the manuscript with support from LF-E, RdH-B, and LT-D. All authors reviewed and approved the final manuscript.

---

**Extended Data Figure 1.**
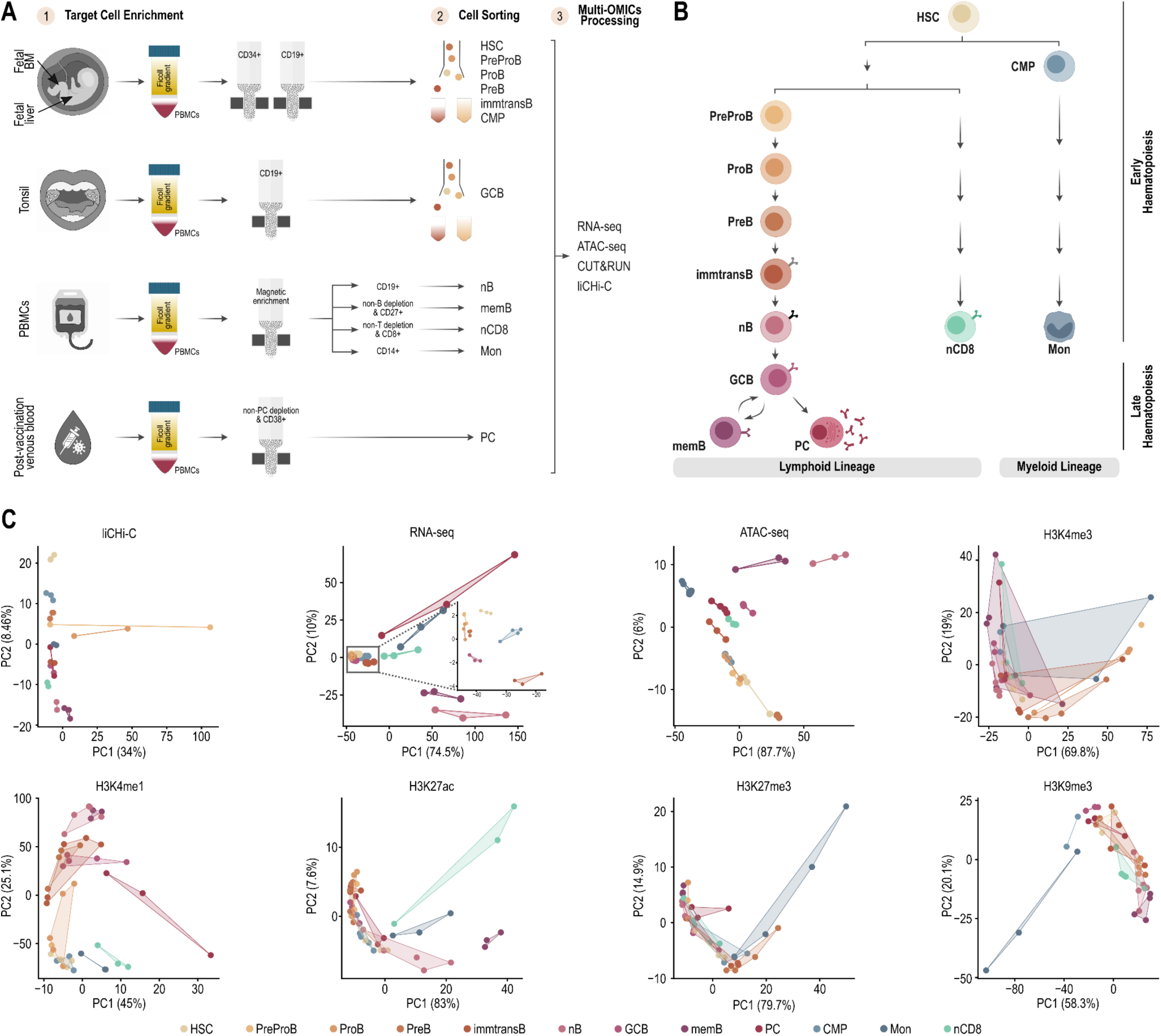
**A.** Experimental framework of cell isolation for multi-omics analysis. **B.** Schematics of hematopoietic differentiation highlighting the 9 B cell-related stages and 3 control cell subtypes used in this study. **C.** Principal component (PC) analysis of liCHi-C significant interactions in at least one sample (CHiCAGO scores >5, n=2-3) (upper row, far left), RNA-seq datasets (n=3-4) (upper row, middle-left), ATAC-seq datasets (n=2-4) (upper row, middle-right), H3K4me3 CUT&RUN datasets (n=1-5) (upper row, far right), H3K4me1 CUT&RUN datasets (n=1-5) (lower row, far left), H3K27ac CUT&RUN datasets (n=2-5) (lower row, middle left), H3K27me3 CUT&RUN datasets (lower row, middle right), H3K9me3 CUT&RUN datasets (lower row, far right).

**Extended Data Figure 2.**
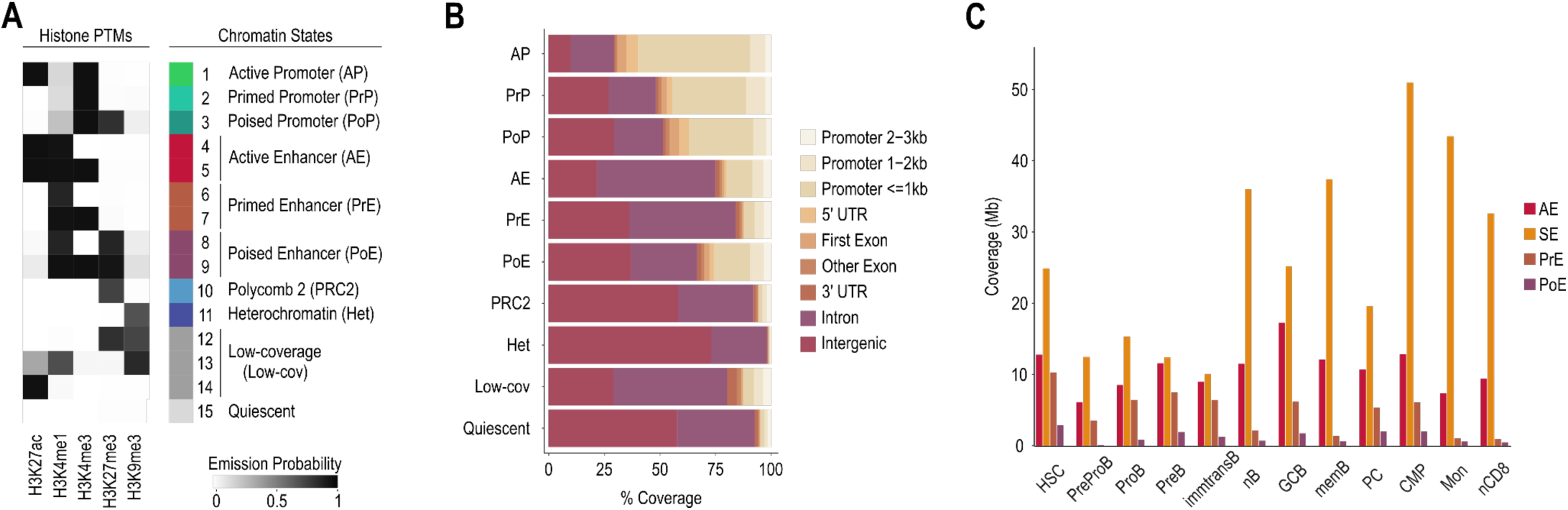
**A.** Heatmap of ChromHMM emission probabilities showing the contribution of individual histone post-translational modifications profiled by CUT&RUN to each inferred chromatin state in a 15-state model. **B.** Genomic distribution of chromatin states, shown as the percentage of genomic coverage across annotated genomic features, including promoters, gene bodies, and intergenic regions. **C.** Total genomic coverage (in megabases) of regulatory element classes - active enhancers (AE), super-enhancers (SE), primed enhancers (PrE) and poised enhancers (PoE) - segregated by cell type.

**Extended Data Figure 3.**
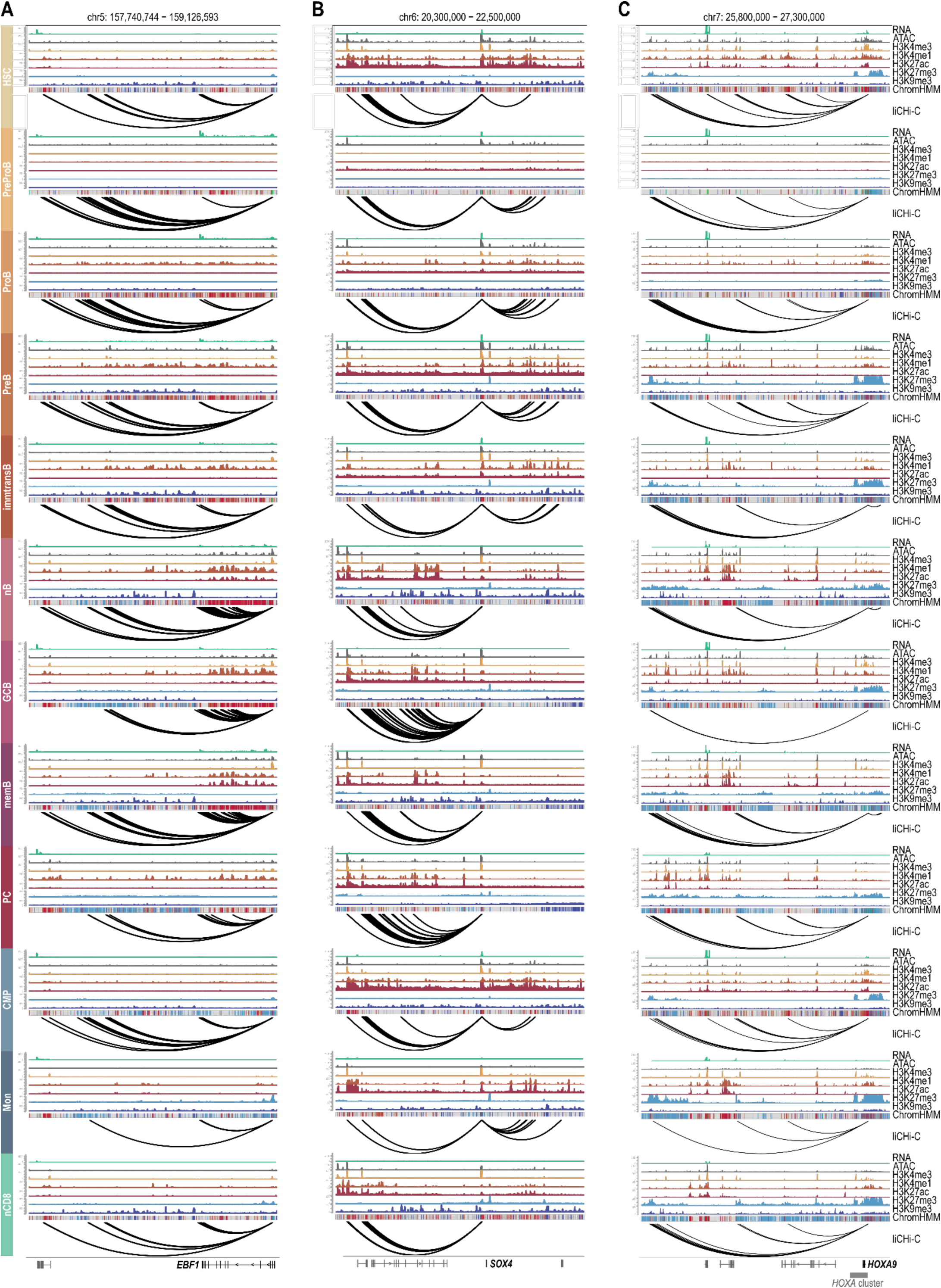
Regulatory landscape in 9 B-cell related cell stages and 3 non-B cell controls, according to RNA, open chromatin, histone modifications, ChromHMM states and liCHi-C data, for *EBF1* (**A**), *SOX4* (**B**) and *HOXA9* (**C**).

**Extended Data Figure 4.**
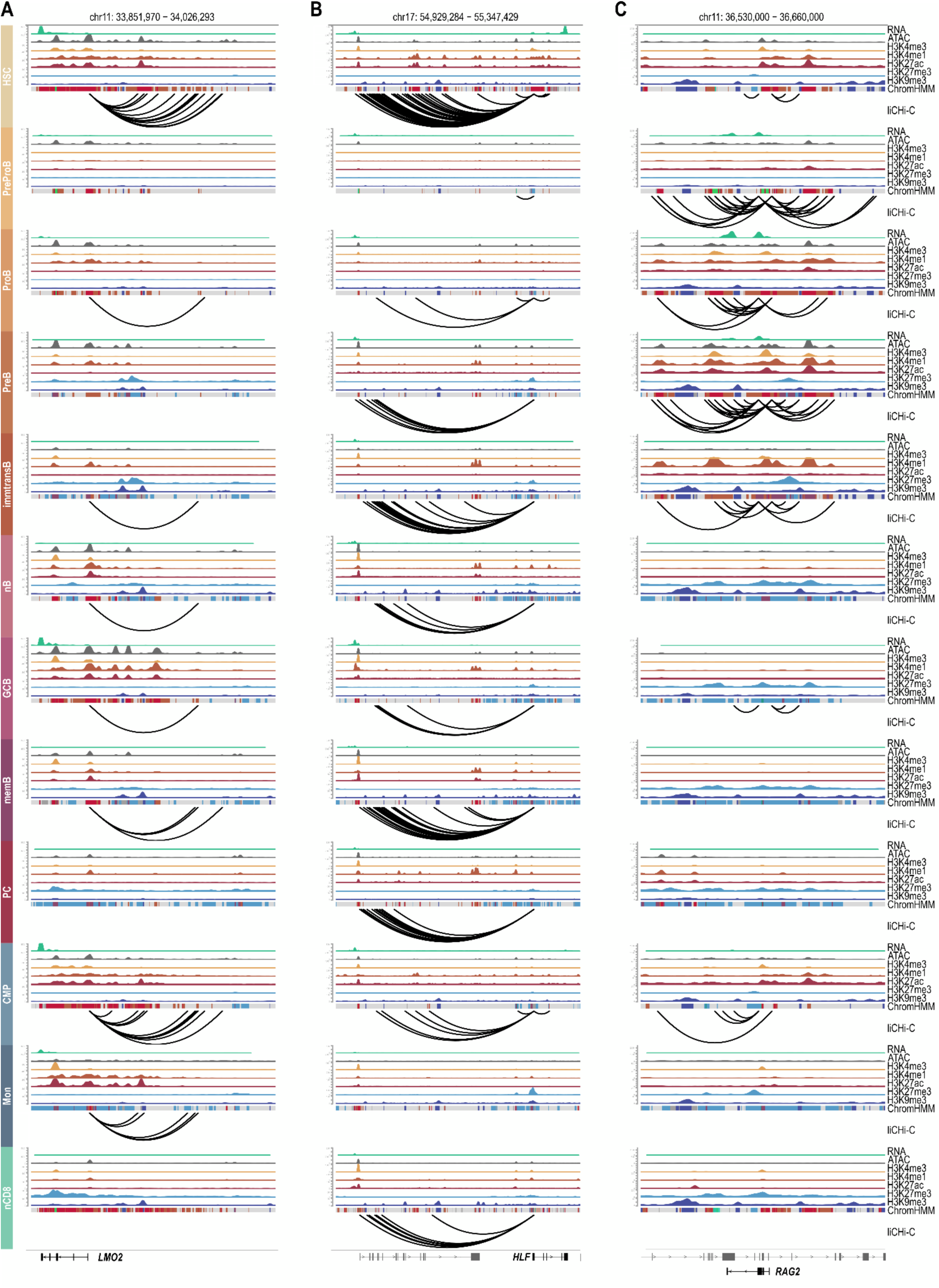
Regulatory landscape in 9 B-cell related cell stages and 3 non-B cell controls, according to RNA, open chromatin, histone modifications, ChromHMM states and liCHi-C data, for *LMO2* (**A**), *HLF* (**B**) and *RAG2* (**C**).

**Extended Data Figure 5.**
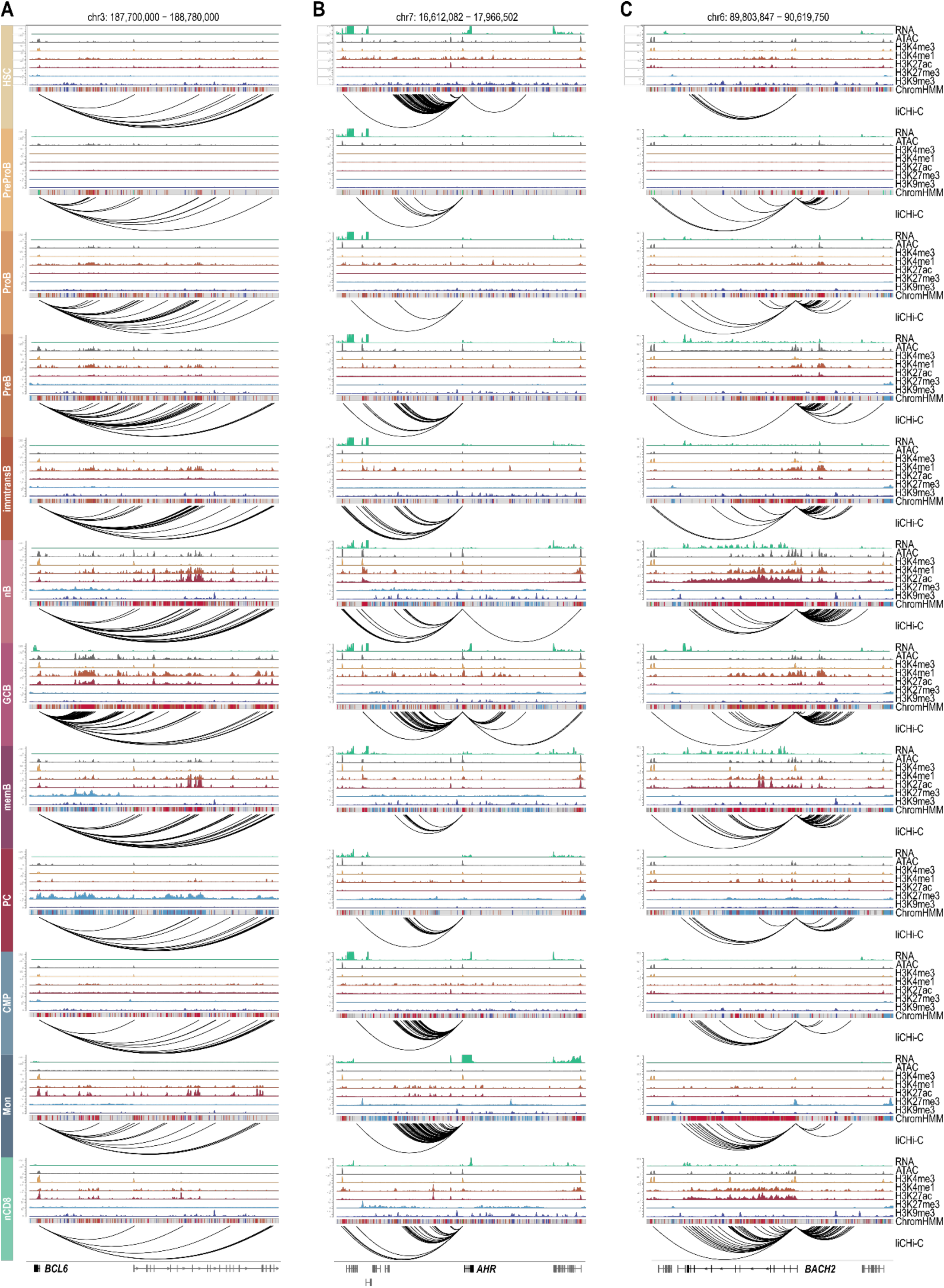
Regulatory landscape in 9 B-cell related cell stages and 3 non-B cell controls, according to RNA, open chromatin, histone modifications, ChromHMM states and liCHi-C data, for *BCL6* (**A**), *AHR* (**B**) and *BACH2* (**C**).

**Extended Data Figure 6.**
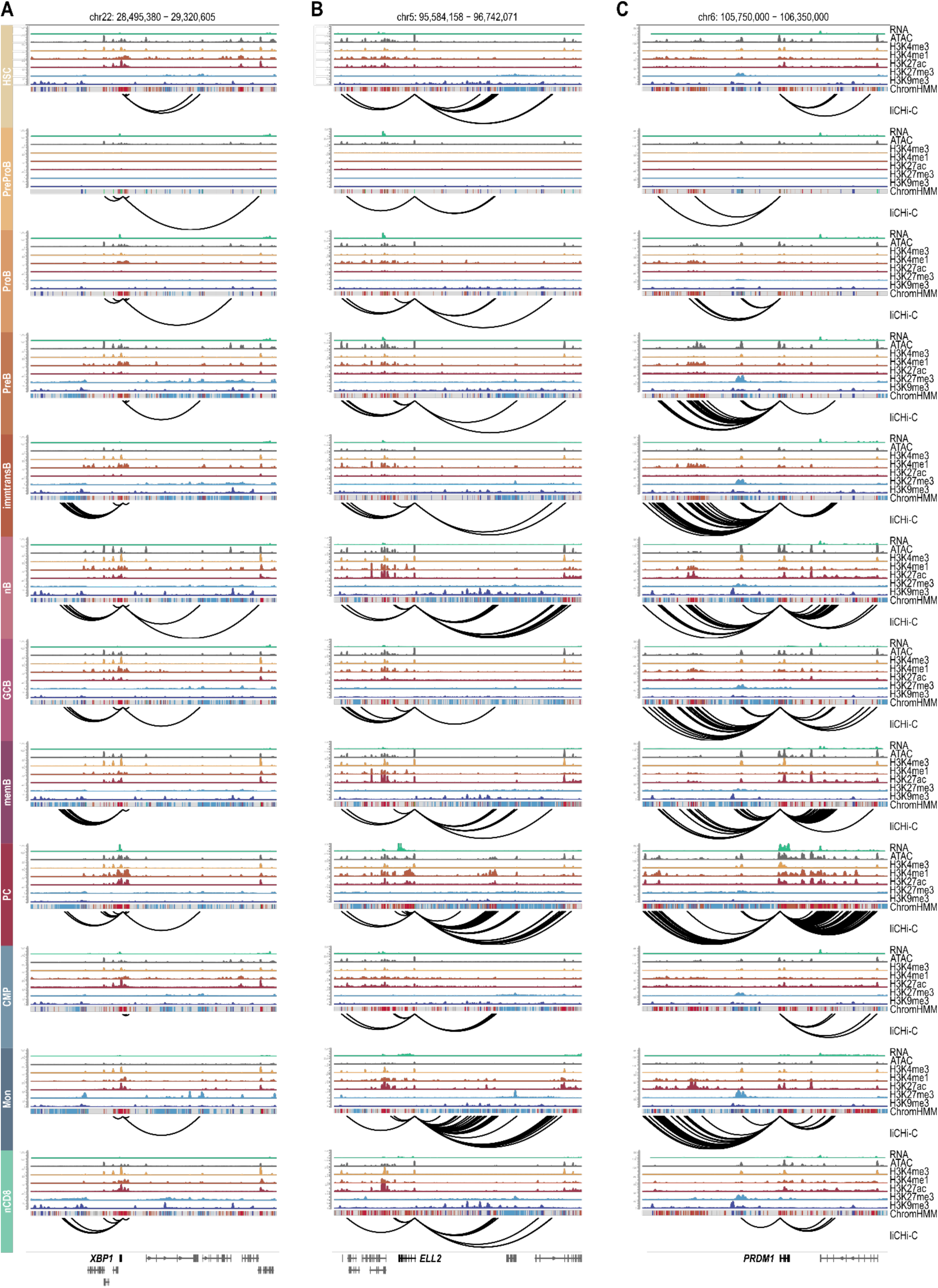
Regulatory landscape in 9 B-cell related cell stages and 3 non-B cell controls, according to RNA, open chromatin, histone modifications, ChromHMM states and liCHi-C data, for *XBP1* (**A**), *ELL2* (**B**) and *PRDM1* (**C**).

**Extended Data Figure 7.**
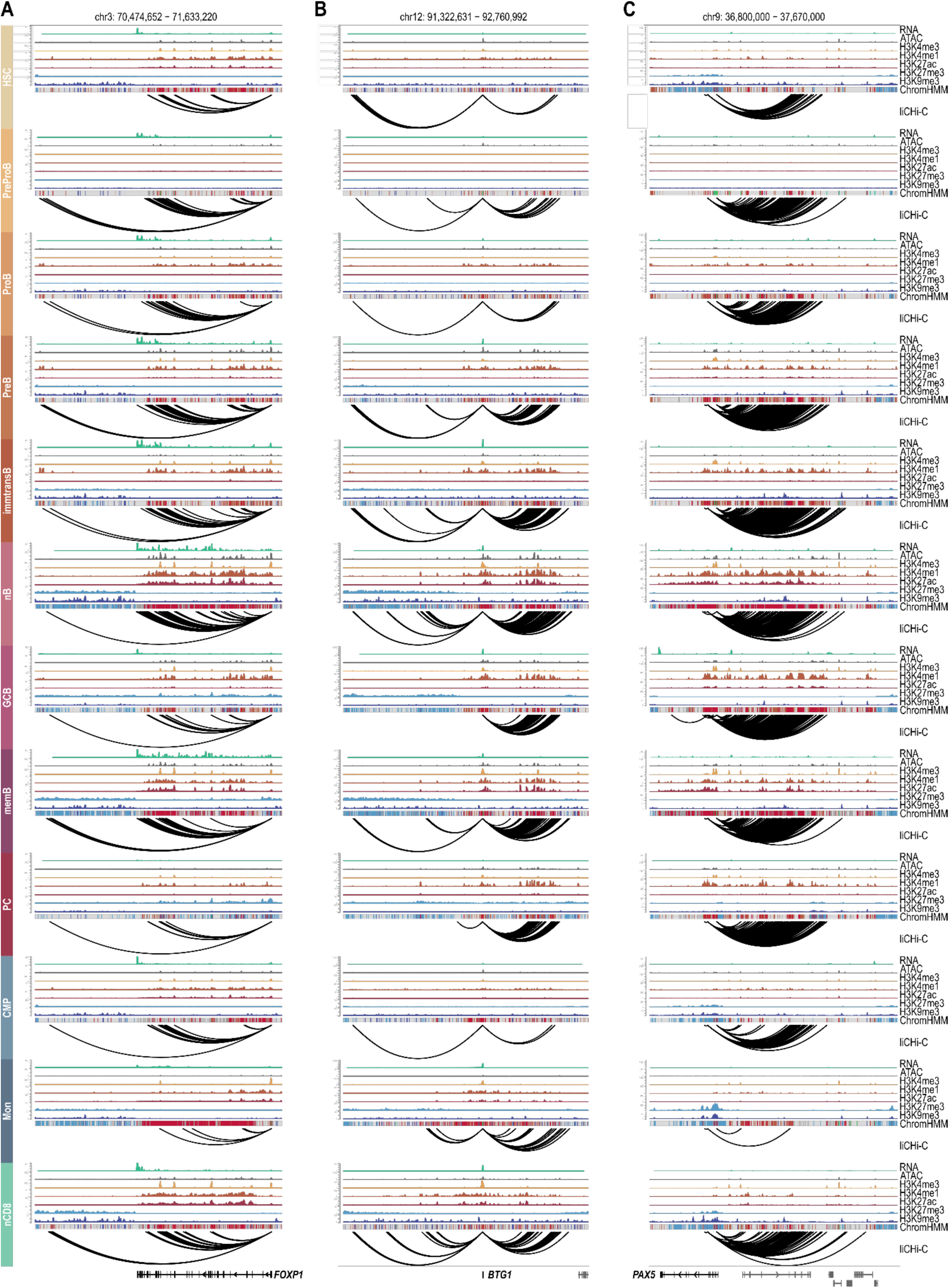
Regulatory landscape in 9 B-cell related cell stages and 3 non-B cell controls, according to RNA, open chromatin, histone modifications, ChromHMM states and liCHi-C data, for *FOXP1* (**A**), *BTG1* (**B**) and *PAX5* (**C**).

**Extended Data Figure 8.**
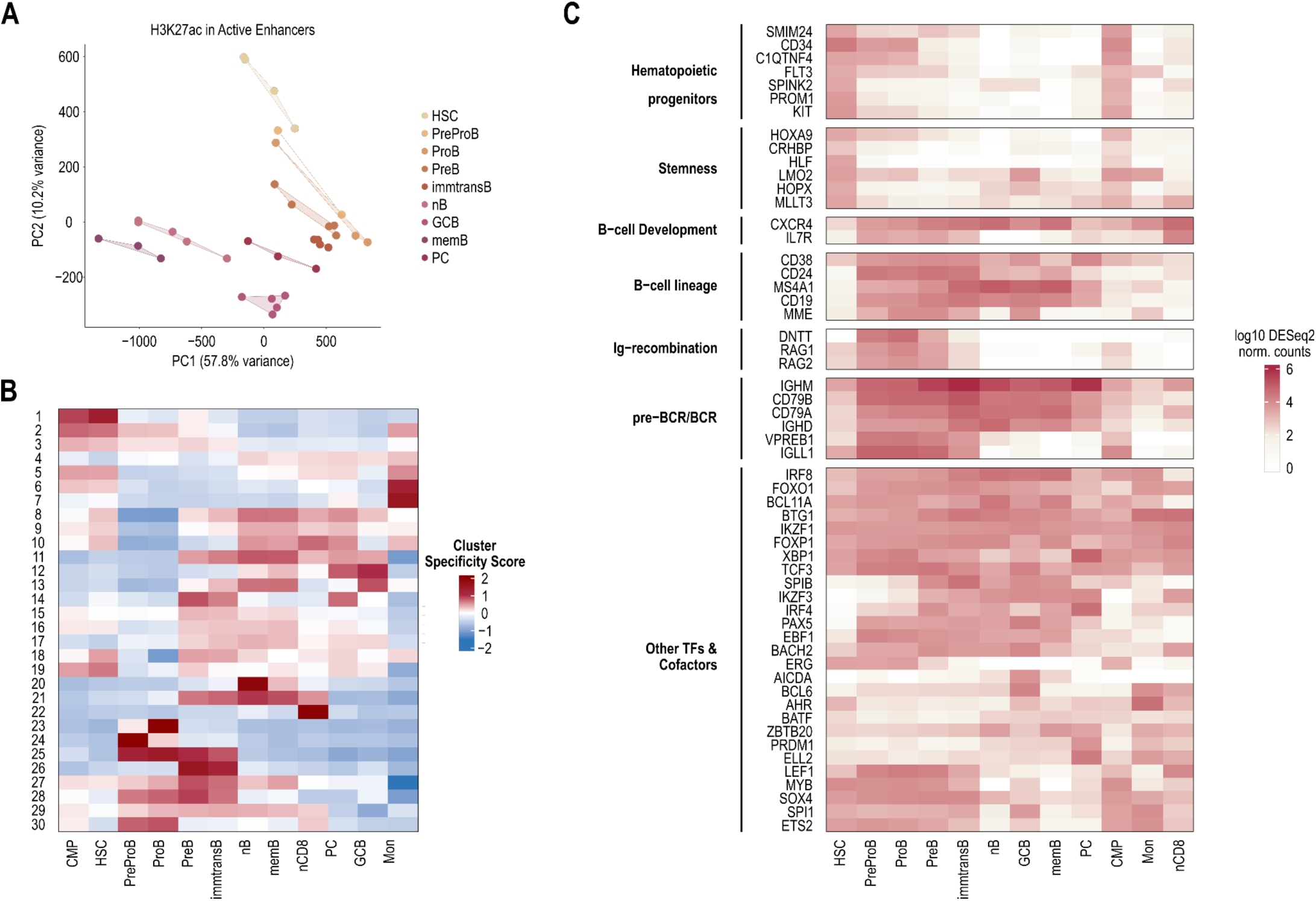
**A.** Principal component analysis (PCA) of H3K27ac signal quantified at promoter-interacting active enhancers across all profiled cell types. **B.** Heatmap showing the cluster specificity score (CSS) for each k-means cluster defined in Figure 2A across cell types. **C.** Heatmap showing the expression of selected B-cell–related genes (including a subset from our manually curated B-gene list) across B-cell differentiation stages and non-B-cell control populations. Expression values correspond to log10-transformed DESeq2-normalized counts, with genes grouped according to functional categories.

**Extended Data Figure 9.**
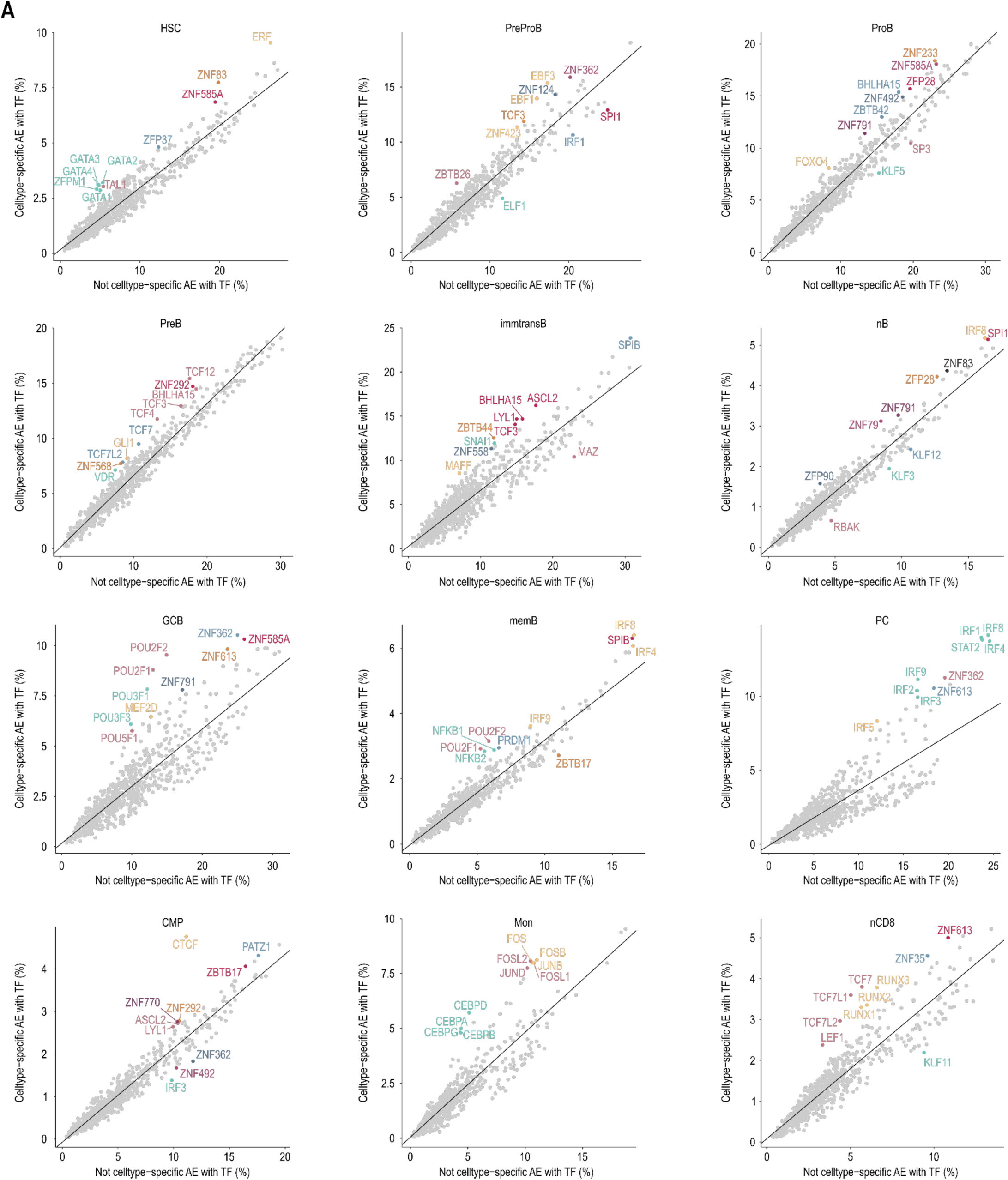
**A.** Transcription factor binding bias in cell-type–specific vs. non-specific active enhancers. Scatter plots showing the percentage of all active enhancers (excluding cell-type-specific) (x-axis) versus cell-type–specific active enhancers (y-axis) bound by each transcription factor (TF) in a given cell type. Each point represents a TF. The diagonal line represents the linear model fit between the two variables. TFs that deviate most strongly (top 10 absolute residuals) are highlighted and labeled. Labeled TFs share color if they belong to the same cluster, defined by motif similarity.

**Extended Data Figure 10.**
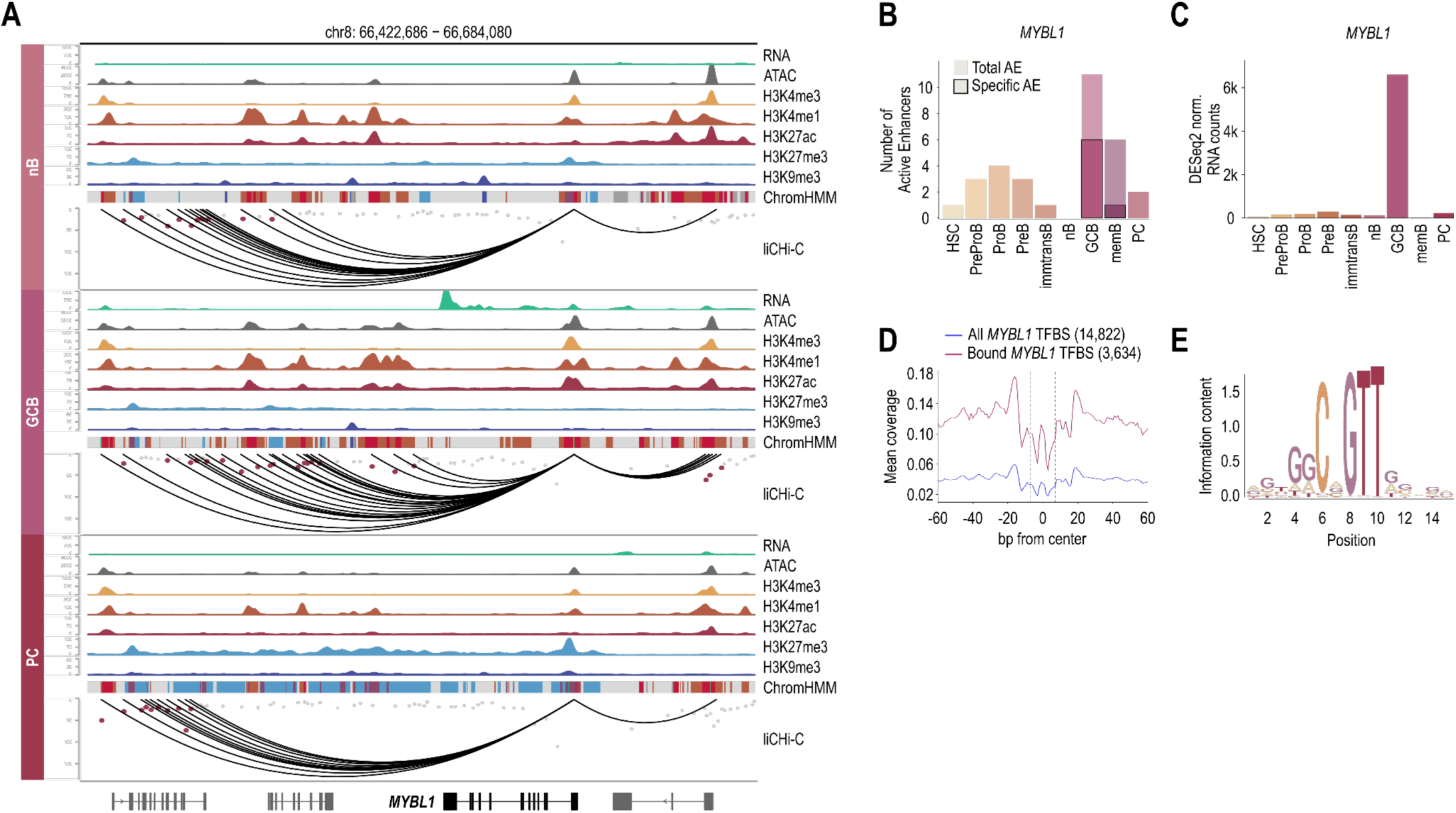
**A.** *MYBL1* regulatory landscape in naïve B cells (nB, top), germinal center B cells (GCB, middle), and plasma cells (PC, bottom) according to RNA, open chromatin, histone modifications, ChromHMM states and liCHi-C data. **B.** Total number of active enhancers (AE) and the subset classified as cell type–specific AE across all profiled cell populations for *MYBL1*. **C.** Gene expression (DESeq2 normalized counts) of *MYBL1* across all profiled cell populations. **D.** Aggregate transcription factor footprinting analysis for *MYBL1* in germinal center B cells (GCB). The blue profile represents the average footprint across all *MYBL1* motif instances overlapping open chromatin regions, whereas the red profile corresponds to the subset of sites predicted to be bound by *MYBL1*. **E.** DNA sequence logo of the *MYBL1* binding motif derived from predicted bound sites. The total height of each position reflects information content relative to a uniform nucleotide distribution, and the height of individual bases represents their relative frequency at that position.

**Extended Data Figure 11.**
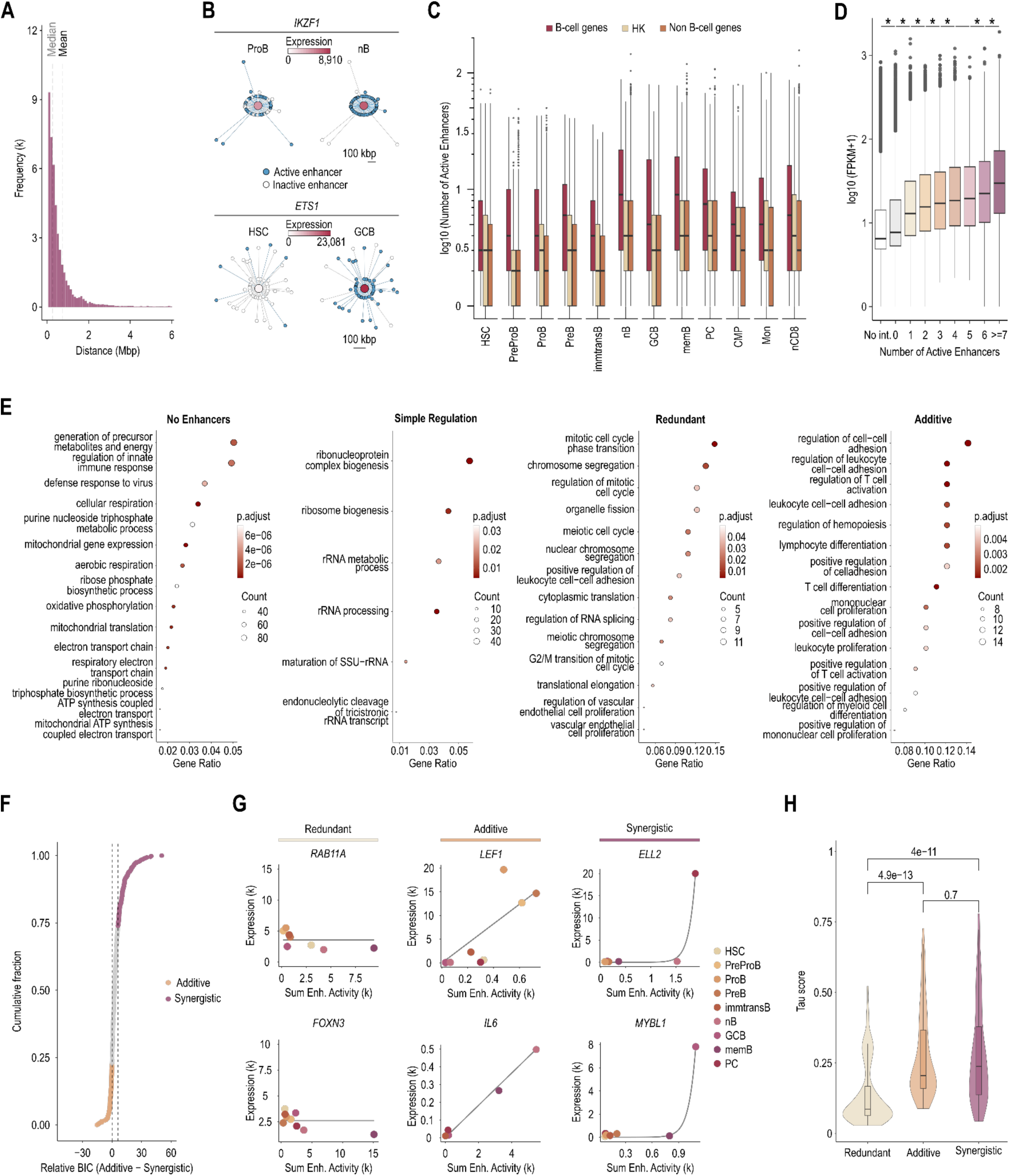
**A.** Frequency histogram showing the distances covered by enhancers within the same gene-centric 3D enhancer network across all analyzed cell types. Distances were calculated as the genomic range between the most distal enhancers linked to the same gene (in megabases). Only networks containing two or more enhancers are included. Grey and black dashed lines indicate the global mean and median distances, respectively. **B.** Gene-centric 3D enhancer networks of *IKZF1* in progenitor B-cells (ProB) and naive B-cells (nB, top); and of *ETS1* in hematopoietic stem cells (HSC) and germinal center B-cells (GCB, bottom). Networks show enhancers that are active in either of the two displayed cell types, arranged radially according to their genomic distance from the target gene (1 unit = 500 kbp) and connected by liCHi-C–detected interactions (grey edges). Enhancers are colored by activity in the indicated cell type (active = blue; inactive = white). Central node color represents *IKZF1* and *ETS1* expression levels in each cell type based on DESeq2-normalized counts. **C.** Boxplot of number of Active Enhancer in each cell type splitted based on the type of genes (B-cell relevant genes, housekeeping genes and rest of the genes). **D.** H3K27ac signal level (FPKM) at promoter of genes stratified by the number of promoter-linked active enhancers, aggregated across all cell types. Asterisks indicate statistically significant differences (Wilcoxon test, p < 0.05) between adjacent groups. **E.** Gene Ontology (GO) enrichment analysis of gene subsets stratified by active enhancer regulation complexity. Dot plots show significantly enriched GO Biological Process terms for genes with no associated active enhancers (No Enhancers), genes regulated by a single active enhancer (Simple regulation), and distal-sophisticated genes regulated by multiple enhancers classified as Redundant or Additive. X-axis indicates the gene ratio for each GO term, dot size represents the number of genes contributing to the term, and color intensity reflects the adjusted p-value. **F.** Cumulative distribution of genes ranked by the relative Bayesian Information Criterion (ΔBIC) between the Additive and Synergistic models (ΔBIC *Add–Syn* = BIC *Additive* − BIC *Synergistic*). Positive values indicate support for the Synergistic model, whereas negative values favor the Additive model. Vertical dashed lines denote the ΔBIC threshold used for model selection (ΔBIC = 6). Genes with ΔBIC *Add–Syn* > 6 and with no support for the Redundant model were classified as Synergistic (purple), whereas genes with negative ΔBIC values against both Synergistic and Redundant models were classified as Additive (orange). Genes with intermediate or conflicting ΔBIC values across model comparisons were considered ambiguous and are not highlighted. **G.** Representative examples of Redundant (*RAB11A*; *FOXN3*), Additive (*LEF1*; *IL6*), and Synergistic (*ELL2*; *MYBL1*) genes. DESeq2-normalized RNA-seq expression is modeled as a function of the summed H3K27ac signal (Enhancer activity) across all interacting AEs for each gene and cell type. **H.** TAU specificity score for the subset of genes classified as Redundant, Additive or Synergistic. Higher TAU values indicates higher cell-type specificity. Statistical differences between groups were assessed using the Wilcoxon test, and corresponding p-values are shown above.

**Extended Data Figure 12.**
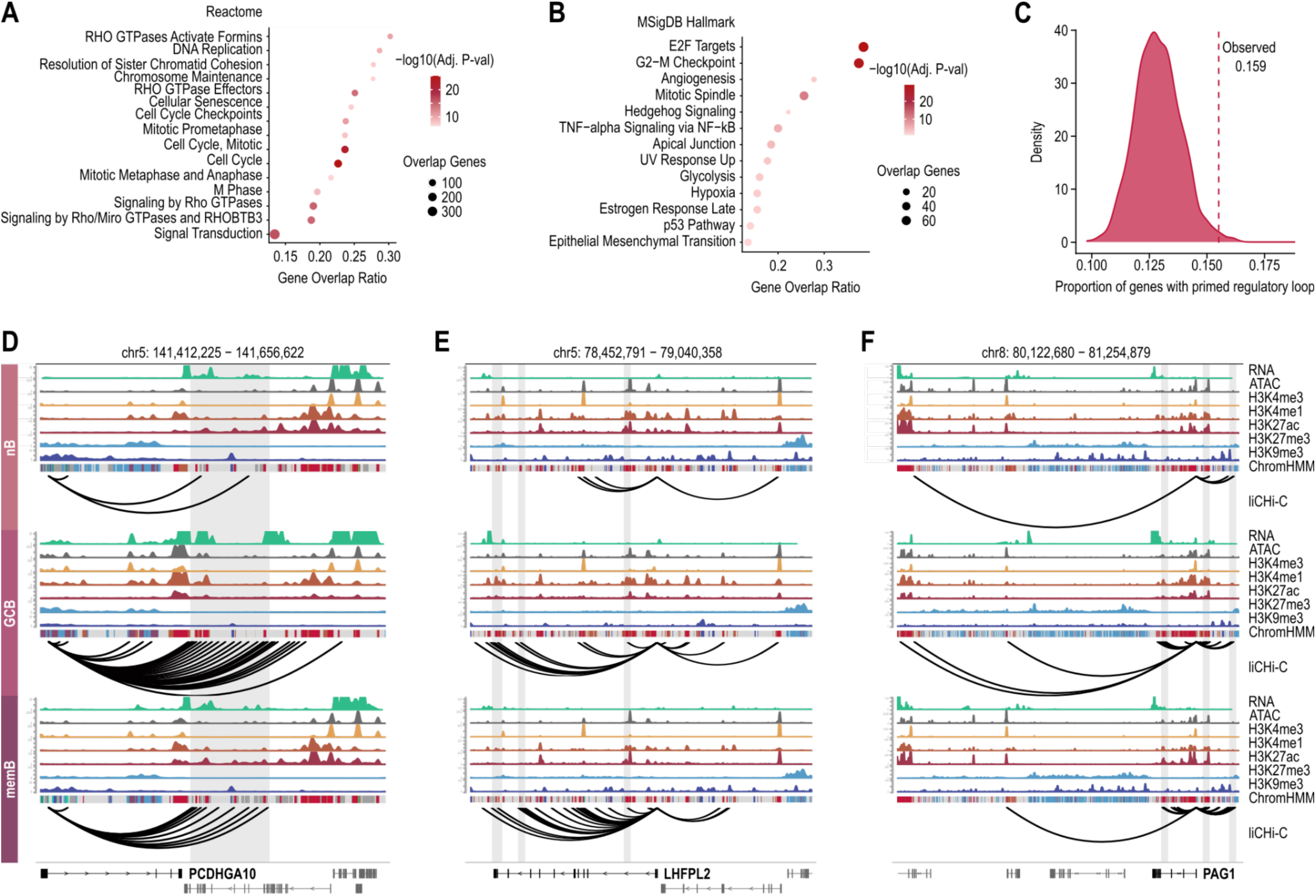
**A.** Pathway enrichment analysis of the 1757 overexpressed genes in germinal center B cells using the Reactome database. **B.** Pathway enrichment analysis of the 1757 overexpressed genes in germinal center B cells using the MSigDB Hallmark database. **C.** Distribution of bootstrapped samples of non-overexpressed genes, showing the percentage of genes with primed regulatory loops. The dashed line indicates the observed percentage for overexpressed genes. **D.** *PCDHGA10* regulatory landscape in naive B cells (nB, top), germinal center B cells (GCB, middle), and memory B cells (memB, bottom) according to RNA, open chromatin, histone modifications, ChromHMM states and liCHi-C data. **E.** *LHFPL2* regulatory landscape in naive B cells (top), germinal center B cells (middle), and memory B cells (bottom) according to RNA, open chromatin, histone modifications, ChromHMM states and liCHi-C data. **F.** *PAG1* regulatory landscape in naïve B cells (top), germinal center B-cells (middle), and memory B cells (bottom) according to RNA, open chromatin, histone modifications, ChromHMM states and liCHi-C data.

**Extended Data Figure 13.**
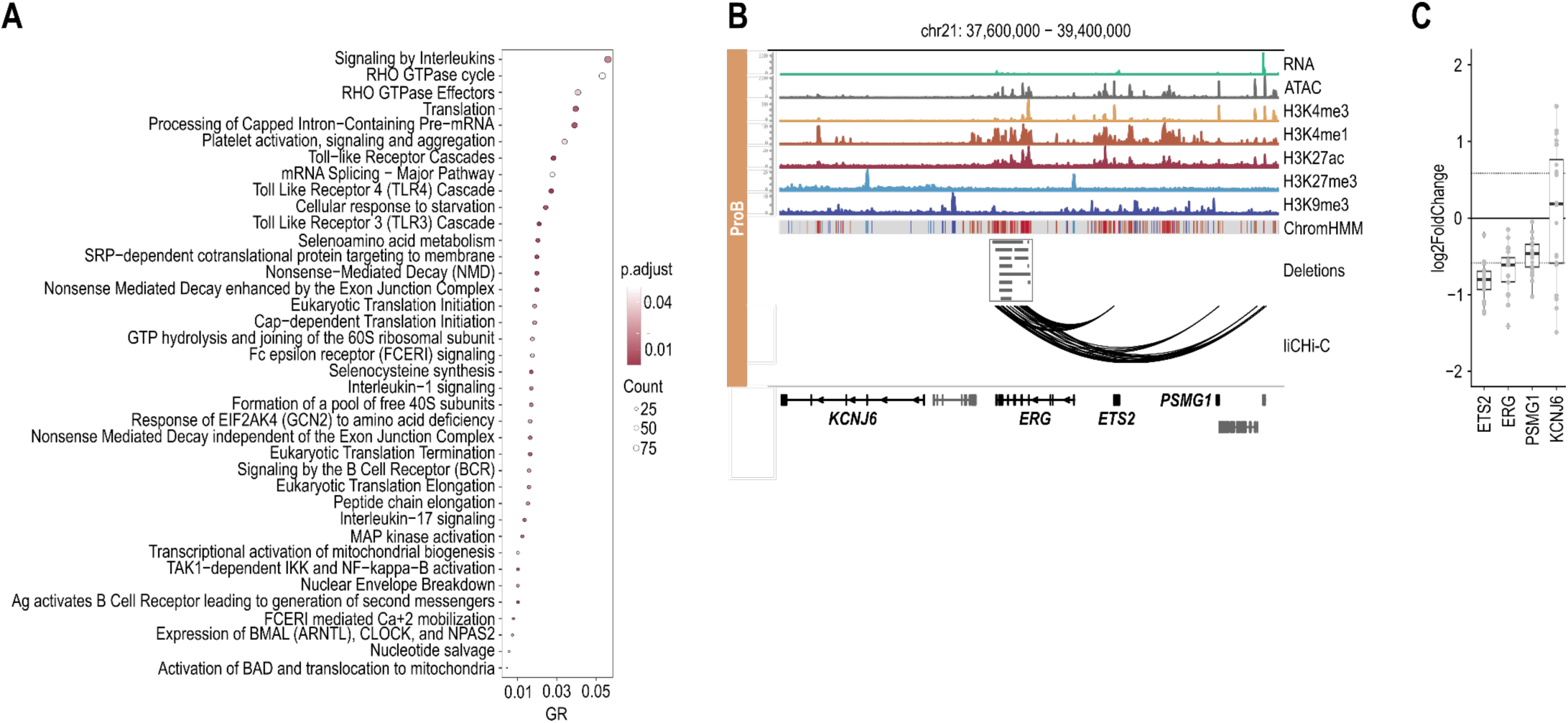
**A.** Reactome pathway enrichment analysis of genes linked to conserved active enhancers in B-ALL patients (light grey/whitish) compared with PreProB-active enhancers. Only pathways exclusively enriched in this gene set are shown, based on equivalent analyses of the other two clusters. **B.** *ERG* regulatory landscape in progenitor B cells (ProB) according to RNA, open chromatin, histone modifications, ChromHMM states and liCHi-C data. The length of the deletions of *ERG* intragenic enhancers across the St. Jude B-ALL cohort are depicted in an independent track (grey shadow). **C.** Log2 fold change in expression of *ERG*, distally interacting protein-coding genes (*ETS2* and *PSMG1*) and nearest but not interacting *KCNJ6* gene, between B-ALL samples carrying deletions affecting *ERG*-linked active enhancers and samples without such deletions in the St. Jude B-ALL cohort.

